# A Constitutive Heterochromatic Region Shapes Genome Organization and Impacts Gene Expression in *Neurospora crassa*

**DOI:** 10.1101/2024.06.07.597955

**Authors:** Andrew T. Reckard, Abhishek Pandeya, Jacob M. Voris, Carlos G. Gonzalez Cruz, Oluwatosin Oluwadare, Andrew D. Klocko

**Author notes:** Equal Contribution (Abhishek Pandeya, Jacob M. Voris). Corresponding Author: Andrew D. Klocko.

## Abstract

**Background:** Organization of the eukaryotic genome is essential for proper function, including gene expression. In metazoans, chromatin loops and Topologically Associated Domains (TADs) organize genes into transcription factories, while chromosomes occupy nuclear territories in which silent heterochromatin is compartmentalized at the nuclear periphery and active euchromatin localizes to the nucleus center. A similar hierarchical organization occurs in the fungus Neurospora crassa where its seven chromosomes form a Rabl conformation typified by heterochromatic centromeres and telomeres independently clustering at the nuclear membrane, while interspersed heterochromatic loci aggregate across Megabases of linear genomic distance to loop chromatin in TAD-like structures. However, the role of individual heterochromatic loci in normal genome organization and function is unknown.

**Results:** We examined the genome organization of a Neurospora strain harboring a ∼47.4 kilobase deletion within a temporarily silent, facultative heterochromatic region, as well as the genome organization of a strain deleted of a 110.6 kilobase permanently silent constitutive heterochromatic region. While the facultative heterochromatin deletion minimally effects local chromatin structure or telomere clustering, the constitutive heterochromatin deletion alters local chromatin structure, the predicted three-dimensional chromosome conformation, and the expression of some genes, which are qualitatively repositioned into the nucleus center, while increasing Hi-C variability.

**Conclusions:** Our work elucidates how an individual constitutive heterochromatic region impacts genome organization and function. Specifically, one silent region indirectly assists in the hierarchical folding of the entire Neurospora genome by aggregating into the “typical” heterochromatin bundle normally observed in wild type nuclei, which may promote normal gene expression by positioning euchromatin in the nucleus center.

## Introduction

Correct genome organization, including proper chromosome folding, is necessary for DNA-templated processes to function normally, including for wild type (WT) control of gene expression [1–5]. Indeed, improper DNA folding altering transcriptional patterns occurs in human cancer cells [6–8]. Eukaryotic genomes hierarchically fold, as DNA molecules wrap around histone octamers to form chromatin fibers, which are extruded into globules (“loops”) by cohesin, and multiple loops compact into Topologically Associated Domains (TADs) [9–17]. TADs of similar transcriptional activity compartmentalize in the nucleus [10, 11, 17]. Specifically, the more open, gene-rich, and actively transcribed euchromatin primarily localizes to the nucleus center, while the silent, gene-poor, and compact heterochromatin is confined to the nuclear periphery [18–20]. Heterochromatin segregation may drive chromatin compartmentalization, although transcriptional activity might force silent genomic loci to the nuclear periphery [19, 21]. In metazoans, the formation of individual chromosome territories – typified by extensive intra-chromosomal contacts – leads to the clustering of heterochromatic regions within chromosomes [22, 23]. In contrast, the independent aggregation of the heterochromatic centromeres and telomeres of fungal chromosomes into a Rabl conformation promotes widespread intra- and inter-chromosomal heterochromatic contacts [9, 24–26]. All told, heterochromatin bundling is a conserved feature of eukaryote genome organization.

Fungi are excellent genome organization models. Fungal genomes are partitioned into euchromatin and heterochromatin, resembling the genomes of metazoans, and the heterochromatin of many filamentous fungi, including *Neurospora crassa*, is subdivided into permanently silent constitutive heterochromatin and temporarily silent facultative heterochromatin [27–29]. The adenine and thymine (AT)-rich, gene-poor, and repetitive constitutive heterochromatic regions, comprising ∼16% of the Neurospora genome, are marked by the tri-methylation of lysine 9 on histone H3 (H3K9me3); this mark is catalyzed by the histone methyltransferase DIM-5 (KMT1) [28, 30–32]. Specific H3K9me3-marked regions found within the seven Neurospora chromosomes (“Linkage Groups” [LGs]) include the regional centromeres, the 14 telomeres, and ∼200 interspersed heterochromatic regions [28, 32]. In contrast, Neurospora facultative heterochromatin is delineated by the subtelomeric di- or tri-methylation of lysine 27 on histone H3 (H3K27me2/3) catalyzed by the SET-7 (KMT6) histone methyltransferase subunit of the Polycomb Repressive Complex 2 (PRC2) [33–35]. In the Neurospora three-dimensional (3D) genome organization, as assessed by in-nucleus chromosome conformation capture with high-throughput sequencing (*in situ* Hi-C), robust intra- and inter-chromosomal heterochromatic contacts occur across Megabases of linear genomic distance, yet heterochromatin minimally interacts with euchromatin, highlighting the compartmentalization of fungal chromatin [9, 24, 25]. Several Ascomycete fungi cluster heterochromatic regions at the base of TAD-like structures, suggesting heterochromatin aggregation may be a conserved mechanism organizing fungal genomes [9, 24–26, 36]. Further, subtelomeric facultative heterochromatin helps chromosome ends associate with the Neurospora nuclear periphery [37]. Together, both heterochromatin types organize the Neurospora genome in a Rabl chromosome conformation.

To understand the importance of individual facultative and constitutive heterochromatic regions in Neurospora 3D genome organization, we performed chromatin-specific *in situ* Hi-C on a strain deleted of ∼47.4 kilobases (kb) within a H3K27me2/3 domain on the LG VI left arm [34], as well as a strain deleted of an ∼110 kb H3K9me3 domain on the LG II right arm (“ΔLGIIK9het25”). While the H3K27me2/3 deletion minimally impacts genome organization, the ΔLGIIK9het25 allele alters the inter-heterochromatic region aggregation, which may increase 3D genome organization variability, and changes the chromatin structure at the deletion site, while nearby AT-rich loci still retain substantial H3K9me3 enrichment. Individual chromosomes in a ΔLGIIK9het25 strain are predicted to have qualitatively different 3D structures than those in a WT strain, and numerous genes are upregulated with the LGIIK9het25 deletion, possibly due to altered gene positioning within the 3D genome. We conclude that a single constitutive heterochromatic region helps form the “typical” genome organization and promotes normal genome function in WT Neurospora.

## Materials and Methods

### Strains and growth conditions

*Neurospora crassa* strains WT N150 (derived from the strain 74-OR23-IVA [Fungal Genetics Stock Center #2489]), N2930 (*mat A his-3;* Δ*mus52::bar^+^*), N3944 (mat x; Δ*dim-5::bar^+^*), and N4933 (*mat a*; Δ47.4 kb::*hph^+^*) were gifts from Eric U. Selker (University of Oregon). All strains were grown with 1x Vogels minimal medium + 1.5% sucrose and supplements at 32°C [38].

The ΔLGIIK9het25 (NKL2) strain, which replaces 110,609 basepairs (bp) of AT-rich DNA in H3K9me3-enriched region number 25 on LG II (between bp 3,602,033 and 3,712,857 of Neurospora reference genome version 14 [nc14]) [24] with 1,401 bp of the *P_trpC_::hph* hygromycin resistance cassette (a net genome deletion of 109,208 bp), was created by split marker gene replacement. Briefly, the ∼1,000 bp upstream of LGIIK9het25 was PCR amplified with oligonucleotides (“oligos”) oKL14 and oKL15, while the ∼1,000 bp downstream of LGIIK9het25 was amplified with oligos oKL18 and oKL19, by Phusion DNA polymerase (cat# F530-L; ThermoFisher Scientific) using the manufacturer’s conditions; oligos oKL15 and oKL18 had ten 5’ complementary nucleotides specific to the *P_trpC_* promoter and *hph* gene 3’ end, respectively. Oligo sequences are provided in Table S1. DNA fragments were gel purified using the GenCatch Gel Extraction Kit (cat# 2260250; Epoch Life Sciences), and split marker fragments fusing the LGIIK9het25 upstream to the *P_trpC_* promoter (“UP”; oKL14 and 2955) or fusing the *hph* gene to the LGIIK9het25 downstream (“DOWN”; oligos 2954 and oKL19) were PCR amplified with the upstream and downstream fragments and a plasmid encoding the *hph* gene (p3xFLAG::hph::LoxP) [39] using LA Taq (cat# RR002M; Takara), and gel purified. The UP and DOWN split marker fragments were transformed into N2930 by electroporation (BioRad MicroPulser Electroporator, cat# 1652100EDU). Hygromycin resistant colonies were selected on 1x Vogels+FGS media with 100 µg/mL hygromycin (cat# ant-hg-5; Invivogen). Genomic DNA was isolated from resultant colonies using standard protocols [40], and tests of proper hygromycin cassette integration at the LGIIK9het25 locus was performed by PCR using oKL49 and 2955 for the left border, 2954 and oKL50 for the right border, and oKL49 and oKL50 for the entire integrated *P_trpC_::hph* gene. Positive integrant strain tKL1A-2 was crossed to N150 (xKL9AR) and genomic DNA was isolated from hygromycin resistant progeny; similar PCR reactions were performed to confirm the presence of the ΔLGIIK9het25 allele in individual cross progeny. PCR with oligos oKL55 and oKL56 determined Δ*mus-52::bar* allele absence, and the strain mating type was established by crosses with known mating type strains. The final homokaryotic strain, NKL2, from cross progeny xKL9AR-17 contains only the ΔLGIIK9het25 deletion (NKL2 genotype: *mat A;* Δ*LGIIK9het25::hph*).

### Hi-C library construction and bioinformatic analyses

*In situ* Hi-C was performed as described [24, 25]. Hi-C libraries were Illumina sequenced at the Genomics and Cell Characterization Core Facility (GC3F; University of Oregon), either on a Hi-Seq 2500 as PE100 reads or a NovaSeq 6000 as PE59 reads. For comparison to mutant Hi-C datasets, WT *Dpn*II and *Mse*I *in situ* Hi-C datasets (NCBI Gene Expresssion Omnibus (GEO) accession number GSE173593) [24] were generated by extracting the number of total reads from WT datasets using the sed command for similar numbers of valid reads as mutant datasets. Merged fastq file reads, or individual replicate reads, were mapped with bowtie2 [41] to nc14 [24], or the nc14-derived reference “NKL2 genome” where the LGIIK9het25 sequence is replaced with the *P_trpC_::hph* sequence. Output sam files were used to build the initial contact probability matrix at a high [1 kb] resolution with hicExplorer software package [42]. Merged contact probability matrices were used to generate lower resolution contact matrices (hicMergeMatrixBins), compare WT and mutant datasets (hicCompareMatrices), KR correct matrices (hicCorrectMatrix), assess TADs (hicFindTADs [using -- correctForMultipleTesting fdr [false discovery rate], --thresholdComparisons 0.3, and --delta 0.15 values for TAD bed files] and hicPlotTADs), and generate images (hicPlotMatrix); replicate similarity was analyzed with (hicCorrelate); additional TAD prediction was performed with Topdom [43]. Chromosome 3D structure prediction was performed with 3DMax (https://github.com/BDM-Lab/3DMax) [44] while Neurospora 3D genome structure prediction was performed by LorDG [45], using custom scripts tailored to fungal chromosome biology; scripts are available on the Oluwadare lab GitHub (https://github.com/OluwadareLab/Fungi_3DGenome). Structure prediction contact matrices were made by merging *Dpn*II and *Mse*I matrices (hicSumMatrices) and converting the .h5 output to NxN text files with a custom script [24] (https://github.com/Klocko-Lab). Output PDB files were displayed on ChimeraX [46] with bins containing differentially expressed genes or chromosomal features manually colored, and high-resolution images were used to create figures. PDB files for all 3D structures are available on Zenodo (<u>10.5281/zenodo.11245552</u>).

### ChIP-seq library construction and bioinformatic analyses

Chromatin Immunoprecipitation-sequencing (ChIP-seq) was performed as described [47], using anti-H3K9me3 (Active Motif; cat# 39161, lot# 30220003) and anti-H3K27me2/3 (Active Motif; cat# 39535, lot# 23120014) antibodies. All ChIP-seq libraries were Illumina sequenced at the GC3F on a NovaSeq 6000 as SR118 reads. Fastq dataset files were mapped to nc14 or the NKL2 genome with bowtie2 [41], converted to bam files, sorted, and indexed with samtools [48], made into bigwig files with Deeptools [49], and displayed on Integrative Genomics Viewer (IGV) [50], with saved png images used for figure creation. Previously published NCBI GEO or the NCBI Sequence Read Archive (SRA) datasets for WT H3K9me3 (GEO accession numbers GSE68897 and GSE98911), WT H3K27me2/3 (GEO accession numbers GSE68897 and GSE100770), Δ*dim-5* H3K27me2/3 (GEO accession number GSE68897 and SRA accession number SRP058573), and input Neurospora DNA (GEO accession number GSE150758) [32, 51–54], were mapped to the nc14 genome [24] and used to create bigwig files, as above, for image generation.

### Total RNA isolation, polyadenylated RNA-seq library construction, and bioinformatic analyses

Total RNA was isolated from overnight liquid cultures of Neurospora strains grown in 1x Vogels, 1.5% sucrose plus supplements [33, 37]. Cultures were harvested by vacuum filtration, placed in 2.0mL screw-top tubes, and 300µL of acid washed glass beads suspended in dH_2_O (150-212µm, Sigma Aldrich cat# G1145-10G), 350µL Acid Phenol:chloroform (5:1; ThermoFisher cat# AM9720), and 350µL NETS (300mM NaCl, 1mM EDTA, 10mM Tris pH 7.5, 0.2% SDS) were added. Samples were vortexed for 10 minutes at maximum speed and centrifuged at 4°C for 5 minutes at 13k rpm. The aqueous layer was transferred to two tubes (∼250µL each), 650µL of cold 100% EtOH was added, and RNA was precipitated on ice for 10 minutes and pelleted by microcentrifugation at 4°C for 10 minutes at 13k rpm. The pellet was washed with cold 70% EtOH (made with DECP-treated dH_2_O), the supernatant was removed, and pellets were allowed to air dry for 15 minutes and resuspended in 50µL DEPC-treated dH_2_O. Insoluble material was removed by centrifugation, and the initial RNA concentration was measured by Nanodrop. Total RNA (10µg) was resuspended in 50µL of DEPC-treated dH_2_O, and the Zymo RNA Clean and Concentrator kit (Zymo Research cat# R1013) was used for RNA cleanup per the manufacturer’s protocol; the 15µL elution was divided into 4µL for Qubit concentration (RNA HS assay kit; ThermoFisher Scientific, cat# Q32852) and TapeStation (Agilent) quality control analysis, while the remaining 11µL was stored at -80oC. Only total RNA samples with high ribosomal RNA to messenger RNA (mRNA) ratios were used for library construction. Polyadenylated mRNA (polyA mRNA) was selected with the NEBNext Poly(A) mRNA Magnetic Isolation Module (New England Biolabs [NEB], cat# E7490L) and polyA mRNA sequencing libraries were generated with the NEBNext Ultra II Directional RNA Library Prep Kit for Illumina (NEB, cat# E7760L and E7765L) per the manufacturer’s protocols; quadruplicate polyA mRNA-seq replicate libraries were constructed for each strain. All polyA mRNA-seq libraries were Illumina sequenced at the GC3F on a NovaSeq 6000 as SR118 reads. Raw fastq files were mapped to the nc14 genome using HiSat2 [55], and htseq-count [56] generated gene count files. Differential Expression analysis between strains was performed with DESeq2 [57] in R/Rstudio [58, 59], selecting significantly changed genes as those with expression log_2_ > 3.0 or log_2_ < -3.0 and an adjusted p-value < 0.001; bed files for display on IGV were generated from the DESeq2 output gene lists in Microsoft Excel, and volcano plots of log_2_ fold change vs. adjusted p-value were generated in R/R studio.

## Results

### Loss of a H3K27me2/3-enriched region minimally affects the local chromatin landscape

Since interactions between regions of similar heterochromatin type comprise many of the strongest intra-chromosomal contacts in the Neurospora genome [24, 25], we examined how deletions of either type, facultative or constitutive heterochromatin, could impact fungal heterochromatin clustering. Starting with facultative heterochromatin, we examined a previously constructed strain, N4933, originally used to assess position-dependent signals for H3K27me2/3 deposition in the Neurospora genome [34]. N4933 harbors a 47.4 kb deletion, from basepairs 175,121 to 222,557 on LG VIL, which removes fourteen H3K27me2/3-enriched genes within a larger facultative heterochromatin domain (this strain is named “ΔK27” for brevity). To assess how ΔK27 impacts genome organization, we performed two *in situ* Hi-C replicates with *Dpn*II to capture chromatin interactions within gene-rich DNA of a higher guanine and cytosine percentage, including in facultative heterochromatic regions. Our replicates show highly similar Hi-C contact matrices across LG VI and are highly correlated (Figure S1), allowing replicate merging into a single Hi-C dataset with 9.2 million (M) valid reads. For comparison, we processed an identical number of valid Hi-C reads from our WT *in situ Dpn*II Hi-C dataset [24]. Our initial observation is that, despite having similar valid read numbers, the ΔK27 dataset is slightly biased in capturing more local contacts and less heterochromatin interactions, likely due to variability in ΔK27 Hi-C experiments, with reduced AT-rich heterochromatin ligation since the underlying DNA has less *Dpn*II sites. Indeed, decreased inter-centromeric contacts in the ΔK27 dataset, relative to the WT dataset, occur in raw contact matrices (Figures 1, S2-S3). Inter-centromeric interactions are present upon Knight-Ruiz (KR) [60] correction (Figure S2-S3), indicating capture of biologically relevant structures. However, the decrease in moderate and long-range contacts in the ΔK27 dataset is evident when this contact matrix is directly compared to the WT dataset (Figure S3). Therefore, we focused on the local chromatin changes on LG VI that manifest with the ΔK27 deletion.

**Figure 1.**
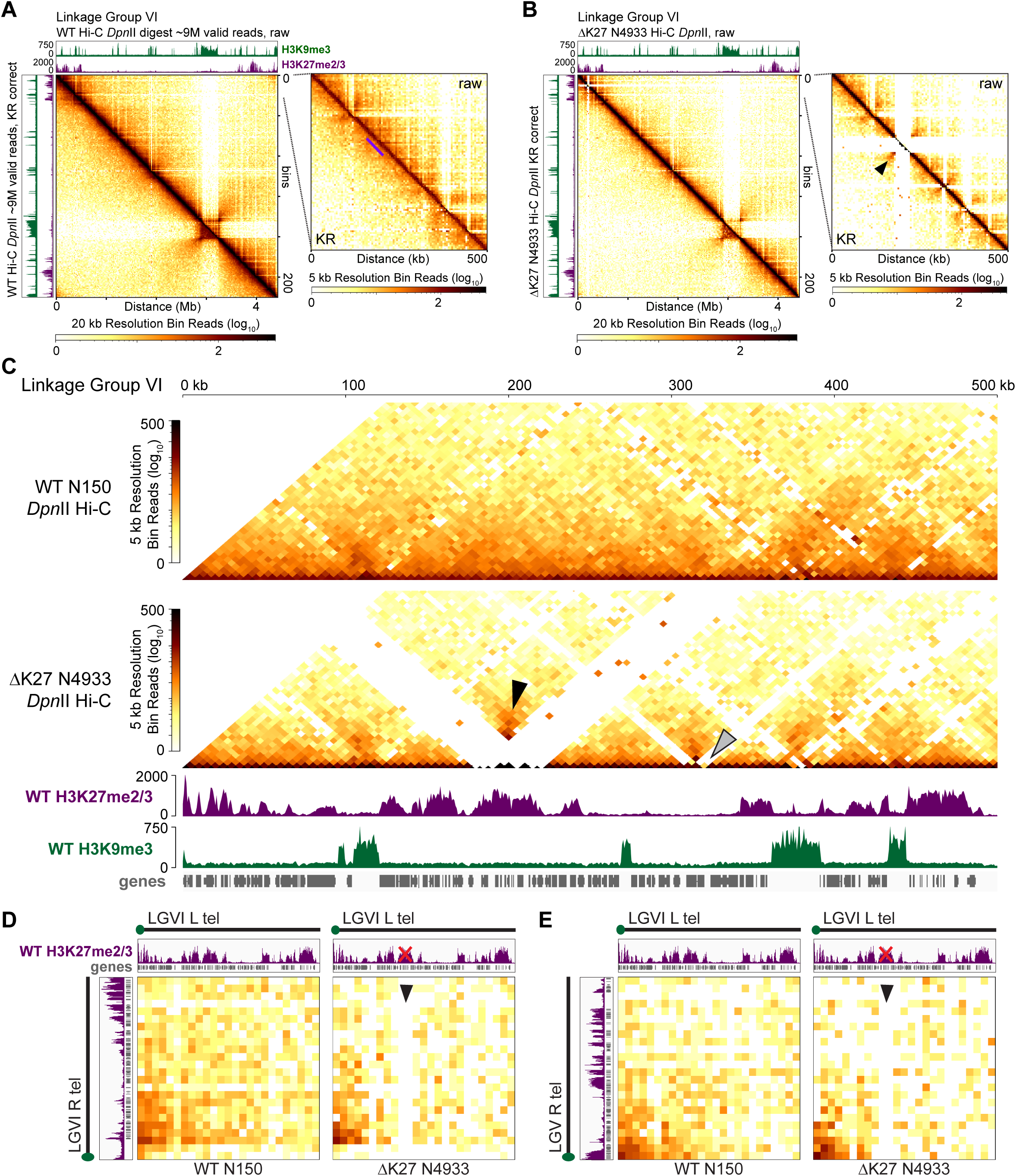
A strain harboring a 47.4 kilobase deletion of a H3K27me2/3-enriched domain has minimal genome organization change. (A-B) Contact probability heatmaps of *Dpn*II (euchromatin-specific) *in situ* Hi-C datasets at 20 kb resolution across Linkage Group VI (LG VI) of the (A) WT or (B) N4933 strains, the latter deleted of a 47.4kb H3K27me2/3-enriched region (“ΔK27”). Enhanced heatmaps show the terminal 500 kb of LG VI, harboring ΔK27, at 5 kb resolution. Each Hi-C image displays the raw contact probability heatmap above the diagonal and the Knight Ruiz (KR) [60] corrected contact probability heatmap below the diagonal; KR correction reduces inherent biases in contact probability matrices. Scalebars shown below the images. Images of wild type H3K9me3 (green) and H3K27me2/3 (purple) ChIP-seq tracks, displayed on IGV [50], above and to the right. The purple line in (A) shows the WT location of the H3K27me2/3-enriched region deleted in the ΔK27 strain, while the black arrowhead in (B) shows the increased contact probability of flanking euchromatin in ΔK27. (C) KR corrected contact probability heatmaps of the *Dpn*II *in situ* Hi-C datasets of WT and ΔK27 strains primarily showing on-diagonal contacts at 5 kb bin resolution. Wild type H3K9me3 ChIP-seq (green), H3K27me2/3 ChIP-seq (purple), and gene (gray) tracks shown below. Scalebar to the left. The black arrowhead shows the increased contact probability of euchromatin flanking the ΔK27 deletion, while the gray arrowhead shows the contact probability loss at a putative transposable element (*sly-1*, gene NCU09969). (D-E) Contact probability heatmaps of the WT and ΔK27 telomere interactions, with the (D) intra-chromosomal (LG VI left and right telomeres) or inter-chromosomal (LG VI left telomere and LG V right telomere) contacts shown. Telomere schematic (with the green circle showing the telomere repeat position), and tracks of H3K27me2/3 ChIP-seq (purple) or genes (gray) shown above and right. The red “X” covers the deleted H3K27me2/3 domain, while the black arrowhead shows the deletion site contact depletion.

On WT LG VI, we observe consistent off-diagonal contacts that decrease with genomic distance, in addition to strong inter-telomeric contacts and the insulation of centromeric DNA from local euchromatin, in both raw and KR corrected heatmaps (Figure 1A). A higher resolution (5kb) WT heatmap of the 500 kb surrounding the region targeted for deletion in N4933 shows a similar inverse relationship between contact strength and genomic distance (Figure 1A). In contrast, the ΔK27 deletion Hi-C, mapped to the Neurospora reference genome version 14 (nc14) [24], appears as a white cross emanating from the deletion locus indicating a gap in mappable Hi-C reads (Figure 1B), since this DNA was not present during Hi-C ligation capture. This gap occurs in both the raw and KR corrected contact probability heatmaps, although the diagonal base in the KR corrected heatmap presents with signal (Figures 1B-C), due to miniscule contact probabilities assigned to bins covering repeated DNA sequences prior to matrix correction (Figure S4); KR correction enhances this bias. Importantly, stronger off diagonal signal at the boundary intersections of the facultative heterochromatin deletion appear (Figures 1B-C, black arrowheads), showing how new interactions occur between the H3K27me2/3-enriched chromatin flanking the ΔK27 deletion. Interestingly, the *sly1-1* transposase gene NCU09969 (Figure 1C, gray arrowhead) is devoid of Hi-C signal in N4933, suggesting the ΔK27 strain had a transposition event. Regarding telomere clustering, the ΔK27 strain still forms intra- and inter-chromosomal subtelomeric contacts (Figures 1D-E), with strong interactions between the terminal ∼100kb of H3K27me2/3-enriched chromatin at chromosome ends. We conclude a deletion of a H3K27me2/3-enriched region minimally impacts the organization of the remaining facultative heterochromatic regions on LGVIL.

### Deletion of a ∼110 kb constitutive heterochromatic region alters silent region clustering

In Neurospora nuclei, gene-poor AT-rich constitutive heterochromatin is compartmentalized from gene-rich euchromatin and most facultative heterochromatin [24, 47]. To ask if an individual constitutive heterochromatic region impacts Neurospora genome organization and function, we deleted a 110,609 bp H3K9me3-enriched region on LG IIR, aptly called “LGIIK9het25” as this region is the 25^th^ constitutive heterochromatic region from the LG II left telomere. We chose to delete this second largest AT-rich region on LG II, since there would be a high probability of observing genome organization changes without compromising chromosome function if the centromere was removed. Using split-marker homologous recombination in a Δ*mus-52* strain [61], we replaced the 110.6 kb LGIIK9het25 with a hygromycin resistance gene (hygromycin phosphotransferase; *hph*) controlled by a strong constitutive *trpC* promoter (a *P_trpC_::hph* cassette; Figure S5A). We note a similar-sized DNA replacement could have more accurately represented the LG II chromosome length, but we wanted to avoid cryptic promoters, repetitive sequences for RIP [28, 62, 63], or AT-rich DNA sequences forming novel constitutive heterochromatic regions within the inserted DNA; progressive LGIIK9het25 region deletions might have elucidated additional changes to heterochromatic loci bundling patterns. We also attempted other heterochromatic region deletions using the nourseothricin acetyltransferase (*nat1*) resistance gene [64], but did not get primary transformants, possibly due to immediate silencing of the AT-rich *nat1* gene at the targeted constitutive heterochromatic loci. We successfully obtained hygromycin-resistant primary transformants deleted of LGIIK9het25 (ΔLGIIK9het25), in which 109,208 bp are removed from the Neurospora genome, given that *P_trpC_::hph* inserts 1,401 bp. A hygromycin-resistant progeny from a sexual cross, xKL9AR-17, was characterized by PCR analysis as harboring both the ΔLGIIK9het25 allele (Figure S5B) and a wild type *mus-52*^+^ allele. The final strain, NKL2 (genotype: *mat A*; Δ*LGIIK9het25::hph*), had no noticeable growth defect from a WT strain (Figure S5C-D).

We performed chromatin-specific Hi-C [24], using either *Dpn*II to monitor gene-rich chromatin contacts or *Mse*I to capture the interactions of AT-rich, gene-poor constitutive heterochromatin. Our ΔLGIIK9het25 Hi-C replicates (four *Dpn*II and two *Mse*I replicates) show highly reproducible and correlated interactions (Figure S6), allowing us to merge replicates into enzyme-specific datasets, with the NKL2 *Dpn*II dataset containing 20.6M valid reads and the NKL2 *Mse*I dataset comprising 12.2M valid reads. All Hi-C datasets display the inter-chromosomal centromeric interactions that occur independently from telomere clusters, indicating capture of biologically relevant structures (Figures S7-S8); these chromosomal features were originally detailed in microscopic observations of fluorescently labeled centromeric and telomeric proteins [25, 37]. As before, we selected similar valid read numbers from previously published WT *Dpn*II and *Mse*I datasets for comparisons [24].

Compared to a WT *Dpn*II Hi-C dataset (Figure 2A), the deleted LGIIK9het25 region appears as a large white “cross” devoid of Hi-C interactions on LGII in the NKL2 *Dpn*II contact probability matrix (Figure 2B). Looking specifically at a 500 kb region surrounding LGIIK9het25, the WT dataset has extensive intra-heterochromatic interactions within LGIIK9het25, and less prominent, but present, heterochromatin-euchromatin contacts (Figure 2A). In contrast, the NKL2 has contact probability loss, as few interactions occur in the raw interaction heatmap and sparse contacts at the diagonal in the KR corrected dataset (Figure 2B), which, like the ΔK27 datasets (Figure S4), is due to a few reads in both the ΔLGIIK9het25 *Dpn*II and MseI datasets mapping to repetitive DNA sequences present in LGIIK9het25, within the nc14 reference genome, increasing contact probability bias (Figure S9). The Hi-C contacts surrounding ΔLGIIK9het25 in NKL2 appear enhanced, as the flanking euchromatin moderately gains contacts in the NKL2 *Dpn*II dataset, relative to a WT dataset (Figure 2A-B). Similar contact probability changes occur in WT and NKL2 *Mse*I datasets, where in the WT dataset the LGIIK9het25 region extensively contacts internal heterochromatin yet is isolated from surrounding euchromatin, but the NKL2 dataset loses intra-LGIIK9het25 heterochromatic contacts yet gains flanking euchromatin interactions (Figure 2C-D). Direct comparison of the WT and NKL2 *Dpn*II and *Mse*I datasets surrounding LGIIK9het25 shows the contact probability gain between flanking euchromatic regions when this separating heterochromatic region is deleted (Figure 2E-F, arrowheads). The surrounding euchromatin may gain local, on-diagonal interactions, as shown in the *Mse*I comparison (Figure 2F), possibly due to either reduced heterochromatic interactions or a local contact capture bias in the NKL2 dataset. Between LG I and LG II, the LGIIK9het25 deletion causes a loss of inter-heterochromatic region contacts, including from the largest non-centromeric heterochromatic region on LG I (Figure 2G, white arrowhead), while the LG II centromere loses inter-centromeric contacts but modestly gains interactions with interspersed heterochromatic regions (Figure 2G, black arrowhead). Similar gains in inter-heterochromatic region interactions are evident across the genome in the NKL2 *Mse*I Hi-C dataset, when compared to the WT *Mse*I dataset (Figure S10), implying a dramatic shift in the heterochromatin bundle with LGIIK9het25 deletion. When comparing the NKL2 *Dpn*II contact matrix to that of a WT strain, the inter-chromosomal contact strength between centromere-proximal euchromatin and more-distant euchromatin on chromosome arms is increased, yet chromosome-internal euchromatic contacts are reduced (Figure S11), consistent with altered bundling between centromeres and interspersed heterochromatic regions. Similarly, both ΔLGIIK9het25 Hi-C datasets have depleted inter-centromeric contacts; KR correction reduces *Dpn*II contact biases (Figures S10-S11). While it is possible that the LGIIK9het25 deletion causes significant genome organization changes, these results may also highlight moderate levels of variability in the Hi-C data derived from the population of nuclei in the NKL2 strain source tissue. Here, valid genome organization differences would be amplified by subtle disparities in Hi-C experiments. These biases occurred despite our use of the same *in situ* Hi-C protocol to generate highly correlated NKL2 replicates (Figure S6), and how we performed the *Dpn*II NKL2 and *Mse*I WT Hi-C experiments at the same time, which hypothetically would minimize experimental bias.

**Figure 2.**
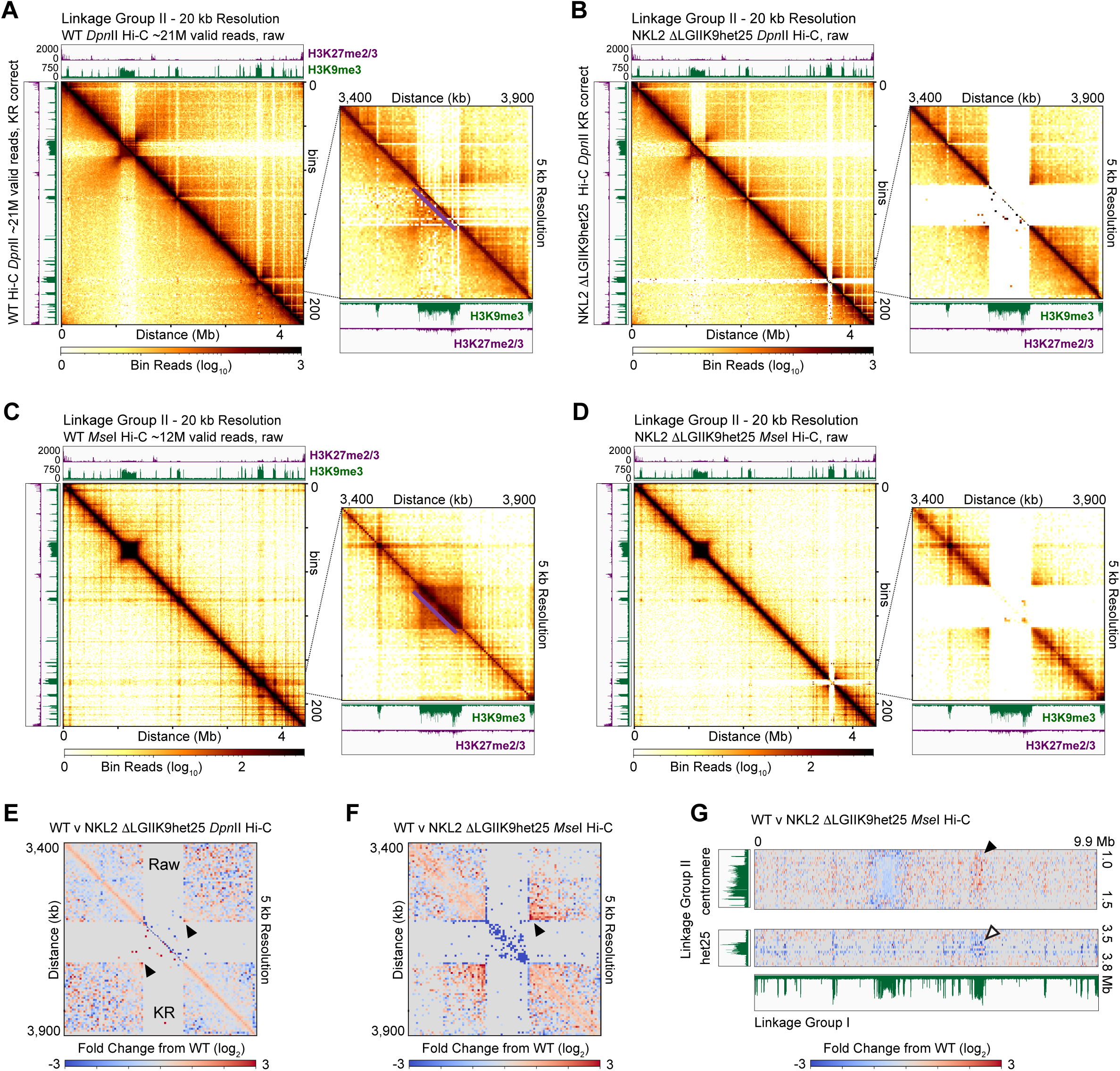
A ∼110 kb H3K9me3-enriched domain deletion alters constitutive heterochromatic region clustering. (A-D) Contact probability heatmaps, presented as in Figure 1, of (A-B) *Dpn*II (euchromatin-specific) and (C-D) *Mse*I (heterochromatin specific) *in situ* Hi-C datasets at 20 kb resolution across LG II of (A, C) WT or (B, D) NKL2 strains, the latter deleted of the 25^th^ H3K9me3-enriched region from the LG II left telomere (“ΔLGIIK9het25”). The purple line in A shows the deleted LGIIK9het25 region. Enhanced (5 kb) resolution heatmaps show the 500 kb surrounding the ΔLGIIK9het25 allele, as in Figure 1A-B. (E-F) Images showing the log_2_ change in Hi-C contact probability between WT and ΔLGIIK9het25 strains in (E) *Dpn*II (euchromatin-specific) or (F) *Mse*I (heterochromatin-specific) 5 kb resolution datasets over the 500 kb surrounding the ΔLGIIK9het25 allele. The scalebar of contact probability changes is shown below. (G) Images showing the change in inter-chromosomal *Mse*I Hi-C contact probability between LG I and the (top) LG II centromere or (bottom) LGIIK9het25 interspersed heterochromatic region.

To assess changes between the WT and NKL2 strains independent of Hi-C restriction enzymes and to compare datasets generated at the same time, we summed the *Dpn*II and *Mse*I datasets from each strain and compared the summed contact probability matrices. The WT and NKL2 summed datasets are similar, with more complete coverage in heterochromatic regions (Figure 12A-B), yet moderately distant contacts in the NKL2 dataset are reduced. Direct comparison of the summed matrices showed the NKL2 dataset with more local contacts and reduction of more distant interactions, with the loss of contacts within the LGIIK9het25 region readily apparent (Figure S12C). Thus, even when some Hi-C experiments performed at the same time are assessed, variation in contact probability, hypothetically due to differences in genome organization that are enhanced in individual Hi-C experiments, is evident. All told, the LGIIK9het25 deletion may affect the organization of the entire Neurospora genome.

### The LGIIK9het25 deletion still allows H3K9me3 deposition at nearby AT-rich loci but alters TAD-like structures

Upon deleting LGIIK9het25, the possibility exists that genome function, specifically H3K9me3 deposition on nearby AT-rich loci, is impacted. To assess if LGIIK9het25 loss prevents histone PTM enrichment on surrounding heterochromatic regions, we performed H3K9me3-specific Chromatin Immunoprecipitation-sequencing (ChIP-seq) on the NKL2 strain. The two NKL2 H3K9me3 ChIP-seq replicates are highly reproducible (Figure S13) for merging into a single file that was normalized by Reads Per Kilobase per Million (RPKM) for direct comparison to previously published WT H3K9me3 ChIP-seq datasets, although using spike-in DNA controls in ChIP-seq experiments performed at the same time may be the best practice for assessing strain differences or relative enrichment levels [65, 66]. We mapped the ChIP-seq reads to either the nc14 WT genome or a “NKL2 genome”, a reference genome in which the *P_trpC_::hph* sequence replaces the AT-rich DNA of LGIIK9het25; enhanced images show low levels of background signal across the *P_trpC_::hph* sequence in an NKL2 H3K9me3 ChIP-seq dataset that are absent in WT H3K9me3 data (Figure S5E).

LG II exhibits an identical pattern of absolute H3K9me3 ChIP-seq enrichment across the AT-rich regions surrounding LGIIK9het25, independent of the reference genome used for mapping reads. For ChIP-seq data mapped to the WT nc14 genome, equal H3K9me3-enriched regions are observed, except peaks to the right of LGIIK9het25 are shifted by ∼110kb from the deletion site (Figure 3A), given that genomic elements positions within our bigwig enrichment count files are shifted based on input bam files having altered read mapping positions across this region. Qualitatively, the NKL2 H3K9me3 peaks may be higher (Figure 3A), possibly reflecting increased H3K9me3 purification from the remaining constitutive heterochromatic regions or enhanced sequencing depth limiting background signal, although we cannot make strict conclusions about relative H3K9me3 enrichment from experiments not performed at the same time. When H3K9me3 ChIP-seq data from both strains was mapped to the NKL2 reference genome, the loss of LGIIK9het25 signal from the WT dataset is apparent, while the remaining constitutive heterochromatic regions have identical LG II genomic positions and absolute enrichment (Figure 3B).

**Figure 3.**
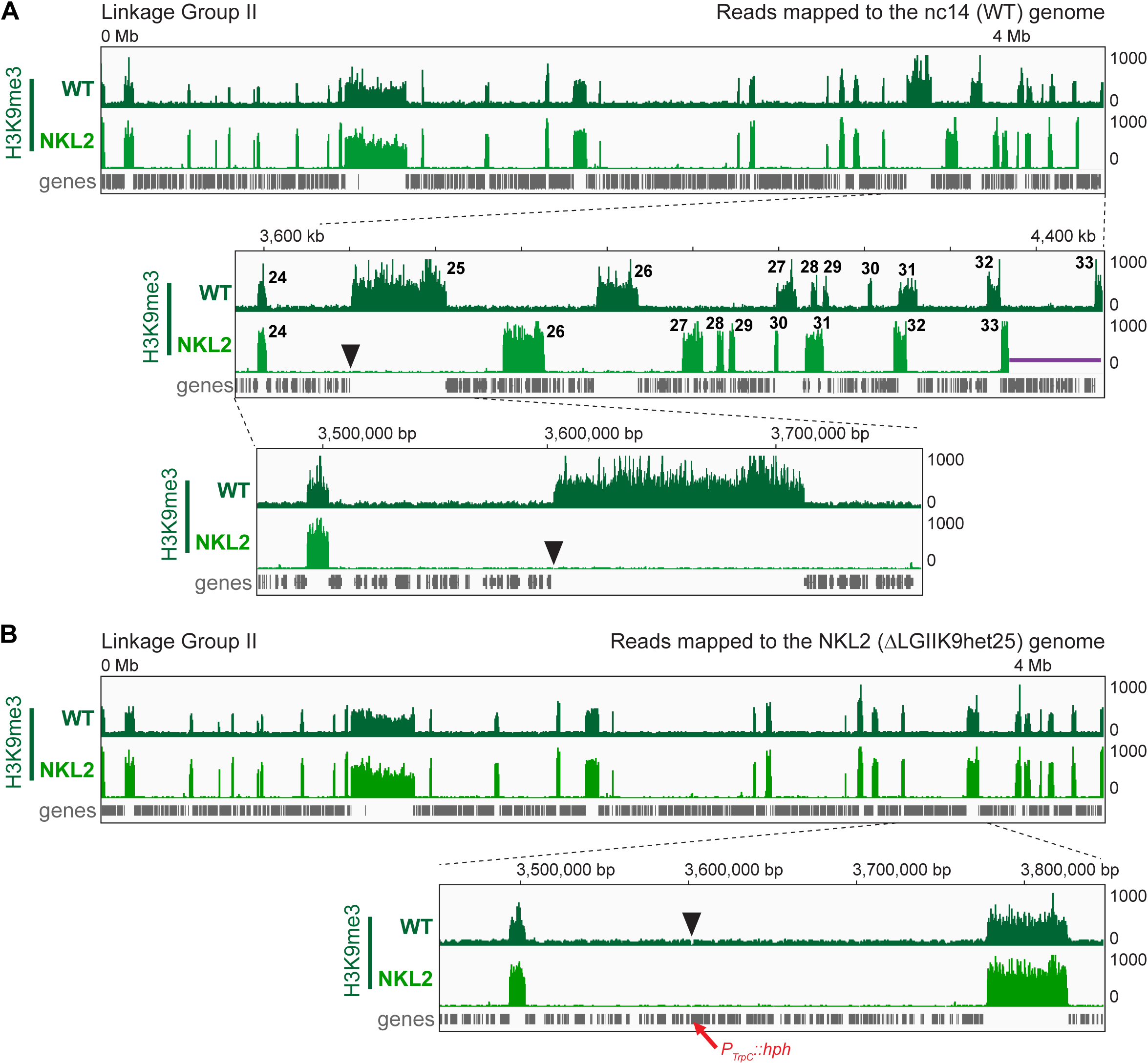
The LGIIK9het25 deletion does not compromise H3K9me3 deposition at other constitutive heterochromatic regions in the NKL2 strain. (A-B) Images of WT (dark green) or NKL2 ΔLGIIK9het25 (light green) H3K9me3 ChIP-seq and gene (gray) tracks, displayed on IGV, of the entire LG II chromosome or enhanced regions. H3K9me3 ChIP-seq reads were mapped to the WT nc14 or (B) NKL2 ΔLGIIK9het25 reference genomes. Numbers in the Figure 3A enhanced panel count the heterochromatic regions from the LG II left telomere, while the black arrowheads show the LGIIK9het25 DNA deletion. The purple line shows the ∼110 kb of DNA missing from the NKL2 genome (shifting all H3K9me3 peaks to the right of the deletion by ∼110 kb). The red arrow in B shows the position of the inserted *P_trpC_::hph* gene in NKL2.

To further assess local chromatin changes resulting from the LGIIK9het25 deletion, we examined TAD-like loop structures on LG IIR. We chose parameters in TAD prediction that mostly placed TAD borders for heterochromatic regions within H3K9me3-enriched domains to increase our chances of predicting biologically relevant TADs, although TAD structure prediction within the same dataset can vary [67]. We started by predicting TAD-like structures in WT and NKL2 *Dpn*II and *Mse*I datasets mapped to the NKL2 reference genome to examine strain-specific changes in chromatin folding without the LGIIK9het25 deletion signal gap. Clear TAD-like structure changes are observed at the deletion site, with the euchromatin in the NKL2 ΔLGIIK9het25 *Dpn*II dataset having larger TAD-like structures, while the NKL2 *Mse*I dataset has smaller on-diagonal TAD-like structures (Figures 4A-B, black arrowheads); this difference likely stems from the chromatin monitored – and the density of restriction sites, with AT-rich heterochromatin having more *Mse*I sites – in each Hi-C experiment. The flanking TAD-like structures are nearly identical in both datasets, except for two larger TADs-like structures in the NKL2 *Mse*I data (Figure 4A-B, open arrowheads). The use of Topdom [43] as second TAD prediction program also showed TAD changes in the NKL2 *Dpn*II Hi-C data relative to WT (Figure S14A), although the TAD changes differ from those predicted by hicExplorer[42]; the Topdom TADs are highly similar in both WT and NKL2 *Mse*I Hi-C datasets (Figure S14B), highlighting the challenges of fungal TAD prediction. Similar TAD-like structure changes across euchromatin are observed in WT and NKL2 *Dpn*II datasets mapped to the nc14 (WT) genome, with larger TAD-like structures predicted with the ΔLGIIK9het25 allele, as opposed to the smaller TAD-like structures in WT datasets (Figure S14C). Interestingly, at the *P_trpC_::hph* insertion, two novel yet small TAD-like structures are observed in both NKL2 datasets, combined with additional stronger Hi-C interactions in *Mse*I data (Figure 4A, purple line), and comparing the NKL2 contact probability and TADs to those in the WT data highlighted the contact probability increase that alters chromatin folding in NKL2 (Figure 4B). These data suggest the constitutive *P_trpC_* promoter could influence the local contacts and expression of genes in this region. Finally, when TADs are predicted in contact probability heatmaps independent of restriction enzyme used (e.g., in summed *Dpn*II and *Mse*I contact probability matrices, mapped to either the nc14 or NKL2 reference genomes), subtle TAD changes are seen across the ΔLGK9het25 deletion site despite good agreement between heterochromatin-euchromatin boundaries and TAD borders (Figures 4C and S14D). Overall, it appears the LGIIK9het25 deletion alters the local chromatin folding surrounding the deletion site.

**Figure 4.**
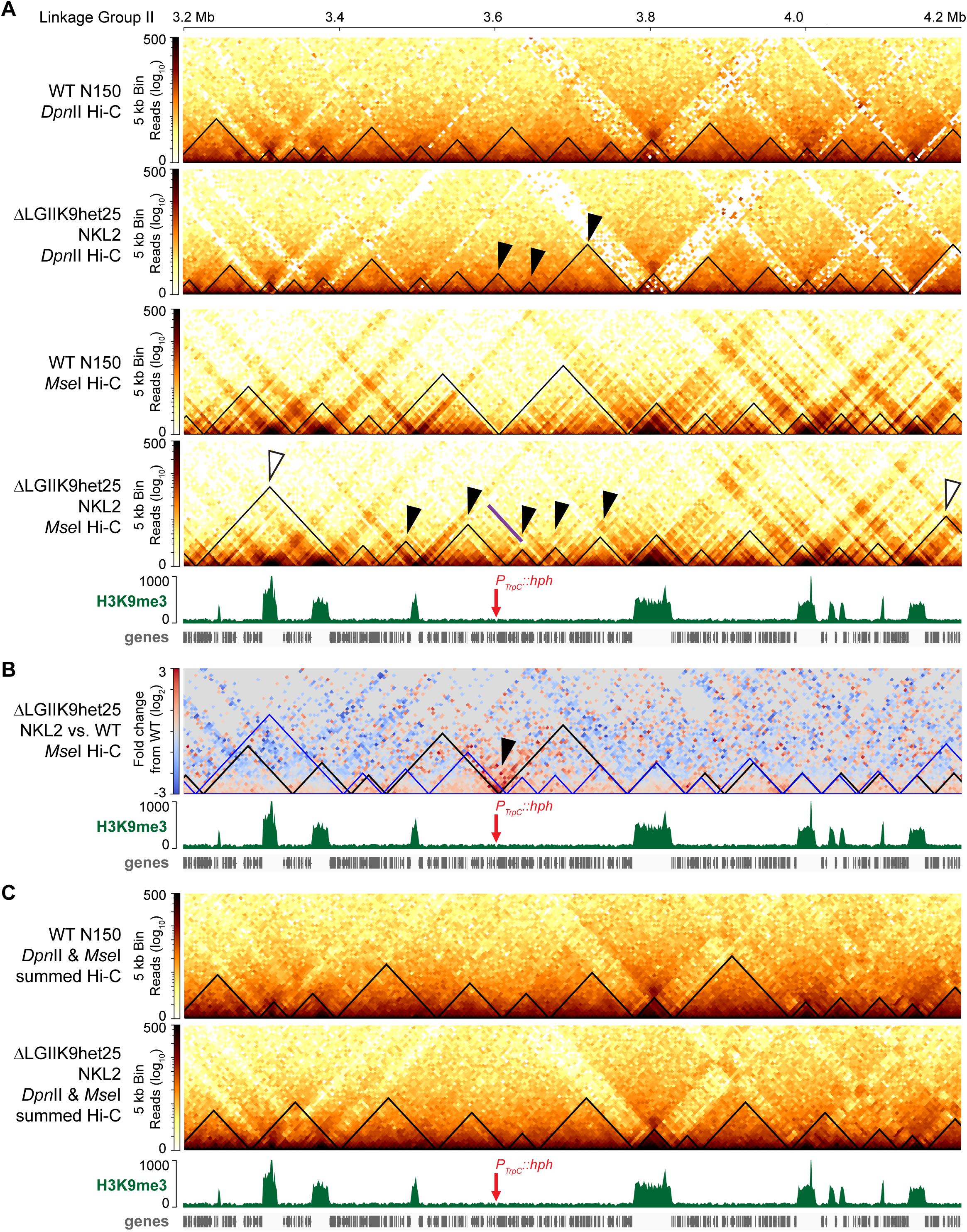
The LGIIK9het25 deletion alters the predicted TAD-like structures in the surrounding chromatin. (A) Contact probability heatmaps of *Dpn*II (KR corrected) or *Mse*I (raw) *in situ* Hi-C datasets of WT and ΔLGIIK9het25 strains primarily showing on-diagonal contacts at 5 kb bin resolution, mapped to the NKL2 ΔLGIIK9het25 reference genome. Wild type H3K9me3 ChIP-seq (green) and gene (gray) tracks shown below. The contact probability scalebar is shown to the left of each heatmap. Predicted TAD-like structures, as determined by hicFindTADs in hicExplorer, are indicated by black triangles. Black arrowheads indicate altered TAD-like structures surrounding the LGIIK9het25 deletion, while white arrowheads show changed TAD-like structures in flanking euchromatin. The red arrow shows the position of the *P_trpC_::hph* gene inserted within the NKL2 reference genome, while the purple line shows increased chromatin contacts at the *P_trpC_* promoter. (A) A heatmap of the changes in contact probability between *Mse*I Hi-C datasets of WT and ΔLGIIK9het25 strains, displayed as in A. The TAD-like structures predicted in WT (black) or ΔLGIIK9het25 (blue) strains by hicFindTADs in hicExplorer are shown (triangles). (C) Contact probability heatmaps, presented as in A, of WT and ΔLGIIK9het25 strains in which *Dpn*II and *Mse*I Hi-C dataset were summed prior to KR correction.

### The LGIIK9het25 deletion alters the three-dimensional folding of Neurospora chromosomes

It remains possible that this large constitutive heterochromatic region deletion alters chromosome folding. To assess the 3D folding of LG II, we computationally predicted the LG II 3D structure using WT and NKL2 ΔLGIIK9het25 Hi-C datasets. We chose to merge all *Dpn*II and *Mse*I data together from each strain to maximize the possible Hi-C contacts across their genomes; we did not combine *Dpn*II and *Mse*I data at ratios equivalent to the Neurospora chromatin distribution, as previously performed [24]. With these summed contact probability matrices, we generated LG II 3D structures using 3DMax [44] customized for biologically relevant structures in fungal genomes; multiple structure prediction iterations occur prior to producing the final 3D structure.

In the WT strain, LG II has an elongated structure with condensed chromosome arms, a more isolated centromere, telomeres localized to a single “side”, and extensive intervening chromatin loops (Figure 5A). Similar 3D structure prediction changes are observed on the other six WT chromosomes (Figure S14). The LGIIK9het25 region forms a loop and associates with nearby chromatin loops (Figure 5A), consistent with nine, moderately sized heterochromatic regions present in the terminal 800 kb of the LG II right arm (Figure 3A). In contrast, the chromosome arms within the LG II 3D structure of the NKL2 LGIIK9het25 deletion are more compact and wider, despite similar centromere isolation, telomere locations, and looping of intervening chromatin (Figure 5B). However, the terminal right arm of the output 3D structure for NKL2 LG II does not form any loops, which suggests that our requirement for maintaining telomere clusters in fungal 3D structure prediction influences how this NKL2 chromosome folds (Figure 5B). Similar folding differences occur on each LG in NKL2, with some chromosomes, particularly LG I and LG IV - having drastically altered 3D structures relative to WT (Figures S14, S15). Together, the LGIIK9het25 deletion modifies the predicted 3D structures of Neurospora LGs.

**Figure 5.**
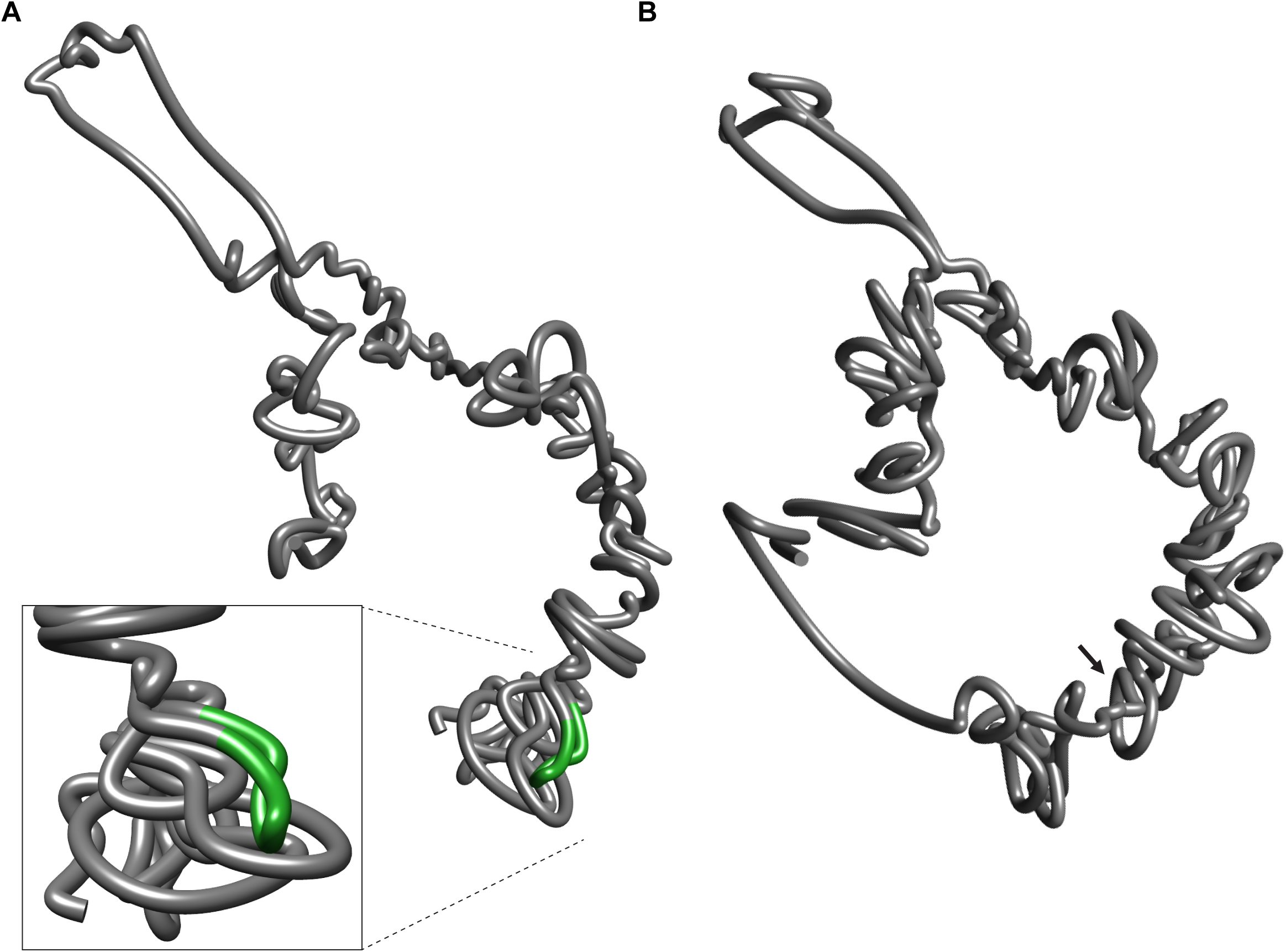
The predicted 3D folding of a single chromosome is changed with the LGIIK9het25 deletion. (A-B) Images depicting the predicted 3D structures of LG II from (A) WT or (B) NKL2 ΔLGIIK9het25 strains at 20 kb resolution. The green region highlights the LGIIK9het25 region in the WT structure. The enhanced image shows the chromatin folding of the LG II right arm surrounding the LGIIK9het25 region in a WT strain. The arrow shows the LGIIK9het25 deletion site.

### The LGIIK9het25 deletion alters gene expression, which is correlated to differences in 3D genome folding

To understand if the removal of LGIIK9het25 could influence genome function by altering gene expression, we assessed changes in polyadenylation messenger RNA sequencing (polyA mRNA-seq) in WT and NKL2 strains. As a control for complete loss of constitutive heterochromatin function, we examined gene expression changes in a Δ*dim-5* strain devoid of H3K9me3 genome-wide [30]. Volcano plots displaying the significant gene expression changes (log_2_ > 3.0 or < -3.0; adjusted p-value < 0.001) in the NKL2 strain relative to the WT strain show 84 genes become significantly upregulated and no genes significantly downregulated, (Figure 6A, left; Supplementary File S1); additional differentially expressed genes (DEGs) could be indicated with less stringent cutoff values. In contrast, the Δ*dim-5* strain has 175 DEGs compared to WT, and many genes have either increased or decreased transcription (Figure 6A, right; Supplementary File S2), including the strong reduction of the gene NCU04402 encoding DIM-5 (Figure 6B), confirming the Δ*dim-5* strain genotype. Many Δ*dim-5* DEGs are found within altered H3K27me2/3 peaks (Figure S16) to possibly explain these transcriptional changes, as facultative heterochromatin relocates to constitutive heterochromatic regions in a Δ*dim-5* strain [51, 52]. The ΔLGIIK9het25 and Δ*dim-5* datasets each have unique DEGs, as only 25 DEGs overlap both datasets. Notably, ΔLGIIK9het25 causes less drastic expression changes relative to the Δ*dim-5* polyA mRNA-seq (Figure 6A-B). Further, no DEGs in the ΔLGIIK9het25 dataset regulate chromatin or transcription (Supplementary File S3), although many DEGs are hypothetical genes without defined functions, preventing a complete tally of the putative regulatory genes. Interestingly, gene expression changes in the ΔLGIIK9het25 strain occur on each chromosome (Figures 6B, S17), with many genes near H3K9me3-marked heterochromatic regions. To assess if linear chromosome distance between DEGs and heterochromatic regions affects gene expression, we measured the closest distance between each DEG transcription start site (TSS) and the nearest heterochromatic region border, plotting the distance in groups (from 1-1,000 bp; 1,001-10,000 bp; 10,001-100,000 bp; and 100,001-1 million [M] bp apart). Most DEGs (n = 51) are 10,001-100,000 bp distant from the nearest H3K9me3-enriched border, with the second most DEGs (n = 23) found up to 1M bp distant (Figure S18), suggesting linear genomic distance has a minor regulatory role. The single gene within 1,000 bp of a heterochromatic region, NCU08696, is directly adjacent to LGIIK9het25 and is downregulated in the Δ*dim-5* strain but upregulated in ΔLGIIK9het25 (Figure 6B, zoomed region), the latter possibly due to its proximity to the strong P*_trpC_* promoter. While longer chromosomes would be expected to have more DEGs, the short ∼4.3 Mb LG VII has more genes with changed expression (n = 24 DEGs) than either the ∼10 Mb LG I (n = 20 DEGs) or the 6.0 Mb LG IV (n =11 DEGs) (Figures 6B and S16, Supplementary File S2). We conclude that a single constitutive heterochromatic region impacts gene expression across the fungal genome.

**Figure 6.**
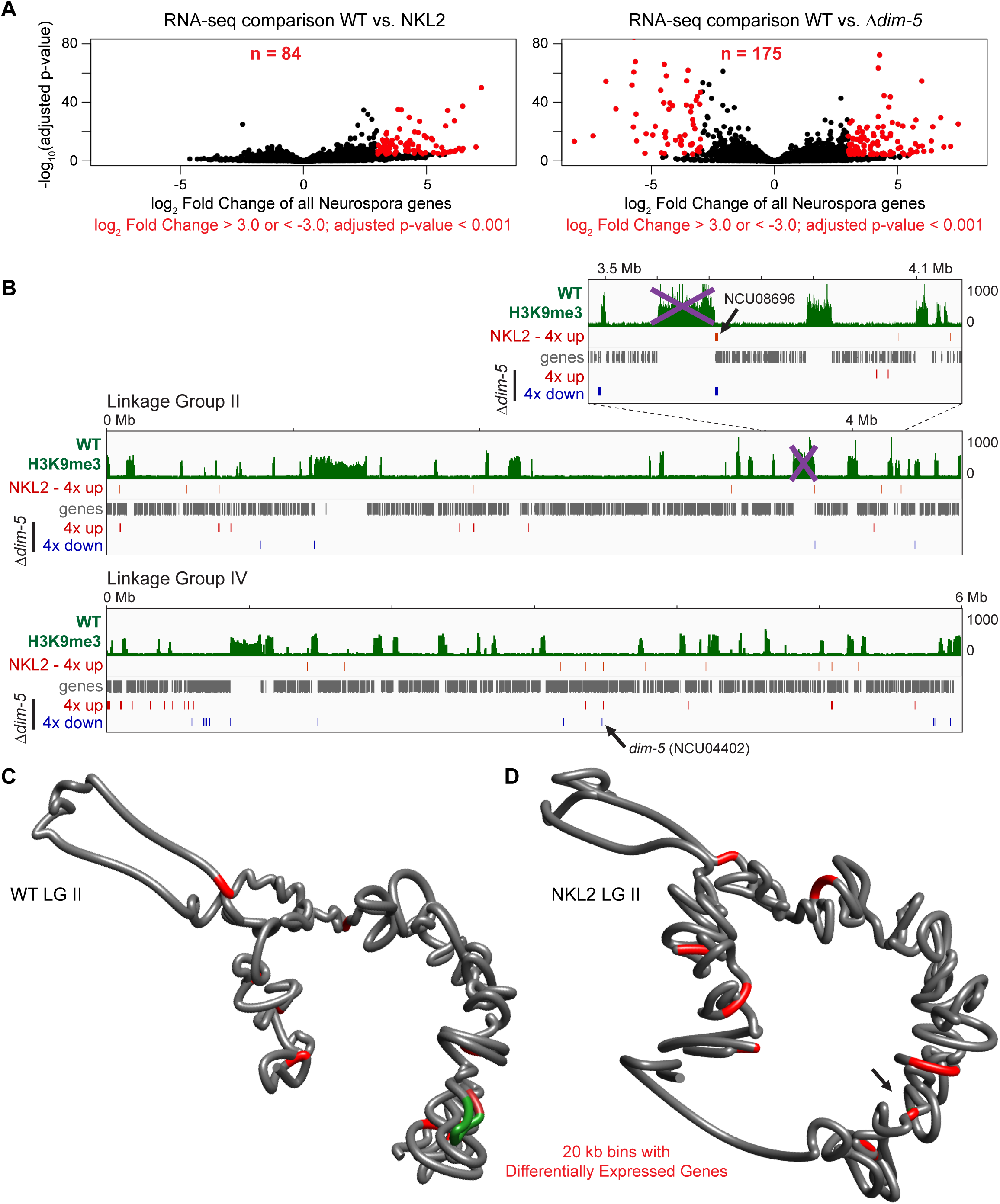
The LGIIK9het25 region deletion causes increased gene expression genome wide. (A) Volcano plots showing the log_2_ fold change from WT expression levels of all Neurospora genes verses adjusted p-values from (left) the NKL2 ΔLGIIK9het25 or (right) the Δ*dim-5* strains. Red points are genes with significant expression changes (log_2_ fold changes > 3.0 or < -3.0 and adjusted p-values < 0.001). The number of differentially expressed genes (DEGs) is indicated at the top. (B) Images of IGV tracks showing WT H3K9me3 ChIP-seq (green), genes (gray), or the “up” (log_2_ > 3.0; red) or “down” (log_2_ < -3.0; blue) DEGs. The purple “X” symbols cover LGIIK9het25 deleted in the NKL2 strain, while the black arrows show the positions of genes NCU08696 or *dim-5* (NCU04402). (C-D) Images depicting the predicted 3D modeling of LG II from (C) WT or (D) NKL2 ΔLGIIK9het25 strains at 20 kb resolution. The green region highlights the LGIIK9het25 region in the WT structure, while 20 kb bins containing DEGs in the NKL2 strain, relative to a WT strain are colored red; the same bins are marked on both structures. The arrow in D shows the LGIIK9het25 deletion site.

To assess if altered 3D chromosome folding could explain the transcriptional changes in the ΔLGIIK9het25 strain, we colored bins harboring DEGs on NKL2 chromosome models. Qualitatively, the positions of the bins containing DEGs are modified on ΔLGIIK9het25 chromosome structures relative to the WT chromosomes (Figures 6C-D, S19, S20). Specifically, on LG II, the bins with DEGs appear more “enclosed” within the WT structure, while these same bins are repositioned to another “face” of the NKL2 LG II structure (Figure 6C-D). Similar qualitative bin movements altering DEG positions are observed on other LGs (Figures S19, S20).

Finally, we examined the positions of ΔLGIIK9het25 DEGs relative to the 3D folding predicted for the entire WT and NKL2 genomes. Our 3D genome models maintain the biologically relevant [25, 37] single centromere cluster which is distinct from telomere bundles (Figure S21). The WT genome model forms an elongated structure with the LGIIK9het25 region close to the structure top (Figure 7A), consistent with the “stretched” conformation of individual chromosomes (Figures 5, S18-S19) and a previous report [68], despite different programs predicting each 3D structure. The NKL2 genome deleted of LGIIK9het25 is wider and less elongated (Figure 7B), similar to the individual NKL2 chromosomes. Intriguingly, the 20 kb bins with ΔLGIIK9het25 DEGs are qualitatively oriented towards the WT genome periphery, while the same bins are more centrally localized in the NKL2 genome, consistent with a hypothesis where enhanced transcription machinery access increases gene expression. We conclude that the deletion of a single heterochromatic region alters the 3D genome conformation and disrupts normal genome function in the Neurospora nucleus.

**Figure 7.**
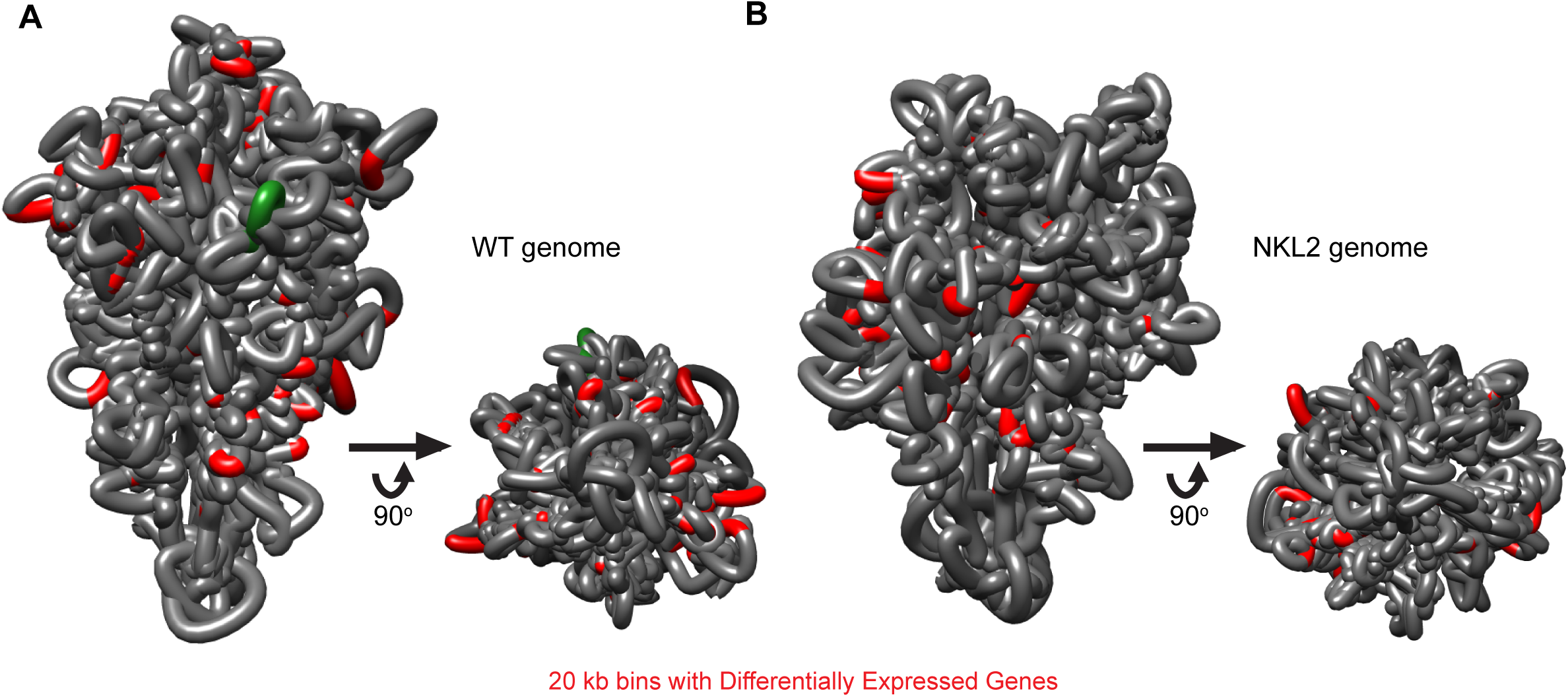
The predicted 3D genome structure and DEG positions are altered upon LGIIK9het25 deletion. (A-B) Images depicting the predicted 3D genome structure of (A) WT or (B) NKL2 ΔLGIIK9het25 strains. The left image is the side view of the genome structure; the right image is where the genome structure is rotated up 90°, showing the centromere cluster. The green region highlights the LGIIK9het25 region in the WT structure, while the 20 kb bins containing DEGs in the NKL2 ΔLGIIK9het25 strain are colored red.

## Discussion

Here, we examined if facultative and constitutive heterochromatic regions organize the fungal genome by performing Hi-C on strains harboring silent region deletions. Our results suggest that gene-rich facultative heterochromatic loci are plastic in nature, as a facultative heterochromatin deletion simply fuses nearby H3K27me2/3-enriched regions within the linear chromosome, which minimally impacts facultative heterochromatin clusters. Specifically, the subtelomere bundling at the terminal ∼150 kb of each LG is maintained in the ΔK27 strain. Subtelomere clustering may be compromised by larger deletions or deletions that target the terminal ∼150 kb of chromosome ends. Consistent with these hypotheses, genome-wide loss of H3K27me2/3 in a Neurospora Δ*set-7* (Δ*kmt-6*) strain can release subtelomere clusters from the nuclear periphery [37], suggesting at least some subtelomeric H3K27me2/3 is needed for telomeres to associate with the nuclear membrane. It is possible that larger or more telomere-proximal deletions would disrupt subtelomere clustering, but further facultative heterochromatin deletions have not been attempted [34]. Still, it is likely that Neurospora tolerates alterations to gene-rich genomic loci, assuming no essential genes are deleted, given the absence of wide-ranging topological or physiological consequences. Since most genes enriched with H3K27me2/3 are novel genes with hypothetical functions [33], perhaps Neurospora tolerates subtelomeric structural variants if facultative heterochromatin is present. In contrast, some Fusarium species require H3K27me2/3 for normal growth [69], meaning structural variants in facultative heterochromatic regions may be lethal in other fungi.

We also assessed the role of a constitutive heterochromatic region for organizing the Neurospora genome. The most recent Neurospora reference genome (nc14) [24] has ∼200 interspersed H3K9me3-enriched regions distributed over seven chromosomes, and apart from the centromeres and telomeres on each LG necessary for chromosome function, the remaining interspersed heterochromatic regions have been dismissed as “junk DNA”. However, when we delete the largest, non-centromeric heterochromatic region on LG II, genome organization and function are impacted without a severe phenotypic effect. Specifically, the heterochromatic regions still present in NKL2 uniquely cluster when LGIIK9het25 is deleted, as we observe gains in intra- and inter-chromosomal heterochromatin contacts, including between interspersed heterochromatic regions and centromeres. Any altered inter-heterochromatic region bundling may reflect valid genome organization changes in NKL2 nuclei, as hypothetically, the LGIIK9het25 deletion would cause the remaining interspersed heterochromatic regions to form novel and variable inter-heterochromatin contacts in each nucleus. These organizational changes may then be amplified by the Hi-C protocol, resulting in the increased contact probability variation we observe. Therefore, a limitation of this study is that we cannot differentiate between these two sources of variability. Still, since genome organization can vary between individual cells across a population [70], it is likely that LGIIK9het25 removal causes contact probability matrix variation from altered heterochromatin bundling. Future controlled fungal Hi-C experiments coupled with computational modeling are warranted to assess the day-to-day Hi-C experiment variability to shed light on valid genome organization changes in WT and ΔLGIIK9het25 strains. Nevertheless, LGIIK9het25 appears to primarily have a structural role in anchoring silent regions on Neurospora chromosomes, while other heterochromatic regions may impact genome function to a greater extent. Additional H3K9me3-enriched region deletions, or progressive truncations of LGIIK9het25, could have elucidated other effects on heterochromatin bundling or fungal genome function. Further, it is possible that additional WT or NKL2 genome organization or function effects could have been discovered with different growth conditions, such as alternative carbon or nitrogen sources, osmolarities, or temperatures, in a manner synonymous to the jet-like chromatin structures across secondary metabolite clusters observed when *Fusarium graminearum* is cultured in excess nitrogen [71].

Our ΔLGIIK9het25 data favor a model where the Neurospora heterochromatin bundle is dynamic, but we cannot exclude the possibility that proteins binding to specific DNA motifs in AT-rich DNA promote heterochromatin aggregation. We suggest that any dynamic heterochromatin clustering could be mediated by Liquid Liquid Phase Separation or a chromatin binding protein, such as Heterochromatin Protein-1 (HP1) [72–74]. Since HP1 loss does not radically disrupt the Neurospora heterochromatin bundle [25], multiple factors, acting redundantly, could drive fungal heterochromatin aggregation. Consistent with this hypothesis, HP1-independent pericentromere clusters are observed in Drosophila [75]. It is currently unknown how eukaryotic constitutive heterochromatic regions bundle, although histone post-translational modifications and/or DNA methylation contribute to fungal genome organization. Specifically in Neurospora, subtelomeric H3K27me2/3 loss causes genome disorder and reduced telomere tethering, while deletion of the HCHC histone deacetylase complex causes constitutive heterochromatin hyperacetylation to increase inter-heterochromatic region contacts; heterochromatin hyperacetylation coupled with 5^m^C loss, reduces heterochromatin compartmentalization [37, 47]. Future experiments should elucidate other chromatin-specific mechanisms for organizing eukaryotic genomes.

The LGIIK9het25 deletion also changes local chromatin structure and 3D genome folding. In a manner synonymous to the ΔK27 strain, the LGIIK9het25 deletion fuses flanking chromatin segments, as shown by contact gains between neighboring euchromatin bins. However, this deletion of LGIIK9het25 alters the predicted TADs, and therefore the regional chromatin looping, across the fused chromatin in two TAD prediction programs. Since interspersed heterochromatic regions in filamentous fungi anchor the bases of TAD-like structures [9, 24–26, 36], in a mechanism tantamount to CTCF forming metazoan loops [10], removing a heterochromatic region could allow interactions between distant euchromatic regions. However, the extensive inter-TAD interactions in fungi [24–26] complicate TAD prediction, and non-uniform TAD-like structures in Neurospora could minimize the functional consequences of altered chromatin loops. While we used TAD prediction parameters to keep heterochromatic region TADs in domains of H3K9me3 to accurately report biologically relevant TADs, one limitation of our study is that euchromatic TAD/loop prediction is particularly challenging in Neurospora. In fact, variable intra- and inter-heterochromatin bundling across chromosomes could cause variability in euchromatin loops in nuclei populations, similar to the variable TAD boundaries reported in different human cell types [76]. Still, our study has functional implications for higher eukaryotic systems. While fungi and metazoans employ some unique chromatin folding mechanisms, the deletion of a metazoan constitutive heterochromatic region could also disrupt TADs to promote new contacts across the normally isolated TAD-internal chromatin, in a manner synonymous to our results in fungi. Thus, loss of a H3K9me3-enriched region from a metazoan genome could alter TAD compartmentalization or misregulate gene expression if several smaller TADs are merged into a larger TAD. Unfortunately, the consequences to metazoan genome function would be more dire, given how TAD boundary dysfunction compromises proper gene expression [6–8]. In fact, our chromosome and genome structure predictions suggest local looping changes could manifest into a unique folding pattern for the entire genome. Here, the ΔLGIIK9het25 deletion caused each chromosome, and the genome, to qualitatively form conformations distinct from the WT genome. While these models are merely predictions and should be interpreted lightly, it is impactful that the repositioning of a single locus has the potential to alter the 3D structure of a whole genome. Hypothetically, despite the species-specific differences in TADs or loops, syntenic changes of linear chromosomal DNA could disorganize a species’ genome, causing functional changes.

Our results also suggest a single constitutive heterochromatic region impacts the transcription of distant genes in the WT Neurospora genome. The gene expression changes we observe in the LGIIK9het25 deletion differ from those in a Δ*dim-5* strain if all H3K9me3 is removed, the latter strain causing numerous genes both strongly up- and down-regulated [37], suggesting LGIIK9het25 has a minor but real impact on genome function. Overall, this single constitutive heterochromatic region may subtly repress Neurospora gene expression. Control of gene expression by H3K9me3-enriched regions in Neurospora was hypothesized from observations of strong contacts between genes and constitutive heterochromatic regions in high-resolution Hi-C datasets [24]. However, it is difficult to imagine that a single LGIIK9het25 region deletion directly causes gene expression changes genome wide. More likely, the ΔLGIIK9het25 allele alters the positioning of the heterochromatin bundle – and the nearby euchromatin – at the nuclear periphery, which repositions some gene promoters for increased transcription machinery accessibility. This hypothesis is supported by modeling the gene expression changes on the ΔLGIIK9het25 chromosomes and across the genome and reasonably explains gene activation in the NKL2 strain. Of course, altered promoters can also drive gene expression changes, as observed by the strong *P_trpC_* promoter in the NKL2 deletion strain increasing transcription of gene NCU08696 immediately adjacent to LGIIK9het25, although the possibility exists that these changes are a consequence of the loss of AT-rich DNA. In contrast, NCU08696, and other genes near constitutive heterochromatic regions, are downregulated in the Δ*dim-5* strain lacking H3K9me3, suggesting that some H3K9me3-enriched regions directly control transcription. Similarly, the proper expression of the Neurospora *met-8* gene requires the immediately adjacent interspersed constitutive heterochromatic region [77]. Together, our results show gene regulation across the fungal genome – even genes on different chromosomes – may depend on a single constitutive heterochromatic region. In metazoans, H3K9me3 enrichment at a gene promoter can repress expression, as the control of mouse embryonic stem cell identity requires the co-deposition of H3K9me3 and H3K36me3 for transcriptional repression [78]. Thus, in addition to the canonical repression of transposable elements and repetitive sequences in eukaryotic genomes [79–82], H3K9me3 can directly regulate transcription. We propose that heterochromatin-mediated gene regulation also occurs through 3D organization, where genes interact with H3K9me3-enriched regions. In this model, genes would associate with constitutive heterochromatic regions at the more repressive periphery for reducing transcription; should these genes be reoriented to the active nuclear center, transcription would increase. It is feasible that constitutive heterochromatic regions could indirectly regulate more “evolutionarily new” or fungal specific genes, in a similar manner as facultative heterochromatin in Neurospora [33], and examining all genes regulated by constitutive heterochromatin in RNA-seq datasets from multiple region deletion strains could assess the evolutionary age of genes controlled by H3K9me3-marked regions. Currently, it is unknown how constitutive heterochromatic regions may regulate gene expression, but an attractive model includes HP1 binding to small amounts of H3K9me3 deposited at gene promoters to allow aggregation with the constitutive heterochromatin bundle; the amount of H3K9me3 needed to implement this regulation is unknown, but could be minor or occur only in a sub-population of nuclei, given the near background levels of H3K9me3 across Neurospora genic regions in ChIP-seq of mycelial populations. Still, fungi may be more apt to employ this hypothetical mechanism to regulate transcription, as opposed to metazoans, given the more-promiscuous contacts across fungal TAD-like structures and the extensive inter-chromosomal contacts in a Rabl chromosome conformation [9, 24–26, 36]. Recently, the histone methyltransferase SETDB1, which catalyzes H3K9me3 in metazoans, was shown to maintain gene expression and drive chromatin compartmentalization [83]. Thus, constitutive heterochromatic regions have additional, underappreciated roles in both genome organization and function across diverse species.

## Conclusions

Our work here has directly assessed how one large H3K9me3-marked silent genomic region impacts fungal genome organization and function. Here, the deletion of a 110.6kb H3K9me3-enriched region alters the hierarchical 3D genome organization in *Neurospora crassa*, with chromatin fibers, TAD-like structures, individual chromosomes, and the fungal genome all having altered conformations. The H3K9me3 deposition on neighboring gene poor regions still occurs despite this 110.6 kb constitutive heterochromatic region deletion, yet this deletion increases gene expression genome wide, which 3D genome modeling suggests is due to upregulated genes being mispositioned into the transcriptionally active nucleus center. Since the Neurospora genome contains 200+ other H3K9me3-regions, it is likely that each silent region similarly contributes to genome function. Since many eukaryotic genomes have H3K9me3-enriched constitutive heterochromatic regions, including many metazoans, the aggregation of constitutive heterochromatin may be a conserved mechanism for organizing eukaryotic genomes and that individual H3K9me3-enriched regions may regulate eukaryotic transcription. Further research in multiple species should help elucidate novel functional roles for this pervasive yet evolutionarily conserved “junk DNA” retained in eukaryotic genomes.

## Declarations

### Ethics approval and consent to participate

Not applicable

### Consent for publication

Not applicable

### Availability of Data and Materials

All genomic datasets are deposited in the NCBI GEO under the SuperSeries accession number GSE269235. Individual datasets have the subseries numbers GSE269228 (ChIP-seq), GSE269229 (Hi-C), and GSE269230 (polyA mRNA-seq). All Neurospora strains are available upon request.

### Authors’ contributions

A.T.R. and A.D.K. designed research; A.T.R., A.P., J.V., C.G.C, O.O. and A.D.K. performed research; A.T.R., A.P., J.V., C.G.C., O.O. and A.D.K. analyzed data; and A.T.R., A.P., J.V., C.G.C., O.O. and A.D.K. wrote the paper.

## Acknowledgements

The authors thank Sara Rodriguez, Yulia Shtanko, and Ashley Scadden for Hi-C library generation help; Sara Hanson (Colorado College) for TapeStation use and RNA-seq protocol advice; the University of Oregon GC3F for Illumina library sequencing service, and Klocko lab members and UCCS Department of Chemistry & Biochemistry colleagues for helpful comments and discussions.

