## Supplemental_Figure-1 for "A Constitutive Heterochromatic Region Shapes Genome Organization and Impacts Gene Expression in *Neurospora crassa*"

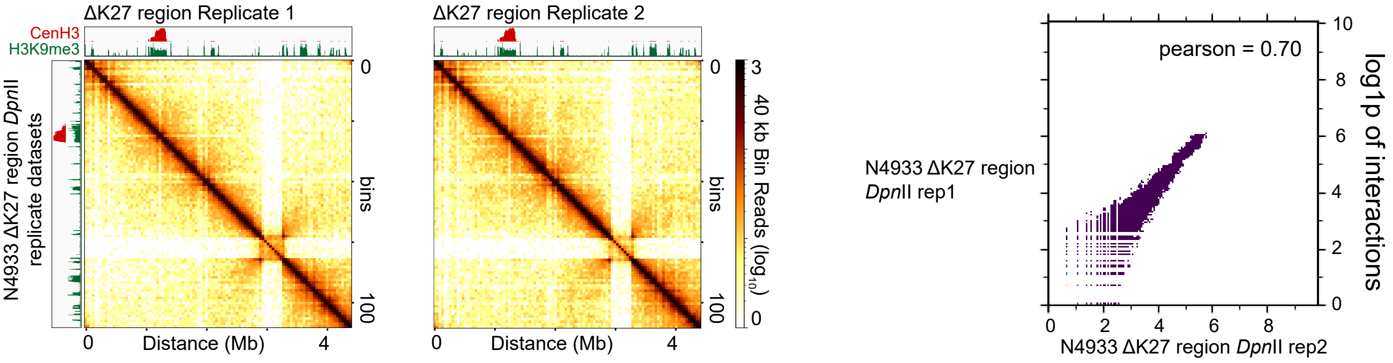


**Figure S1. Replicate analysis of *in situ* Hi-C genome organization datasets, using the *Dpn*II restriction enzyme, of the ΔK27 region strain of *Neurospora crassa*.** Heatmaps and scatterplots of ΔK27 region replicates digested with *Dpn*II. In each panel, the left images show the raw count Hi-C heatmaps of genomic interactions across LG VI of each *Dpn*II-derived *in situ* Hi-C replicate dataset at 40 kb bin resolution. The right image shows a scatter plot comparing interactions between each ΔK27 region DpnII replicate matrix at 20kb bin resolution; log1p values of interactions presented. Pearson correlation values of each binary comparison shown at the upper right. Image produced by the hicCorrelate program in hicExplorer.
