## Supplemental_Figure-2 for "A Constitutive Heterochromatic Region Shapes Genome Organization and Impacts Gene Expression in *Neurospora crassa*"

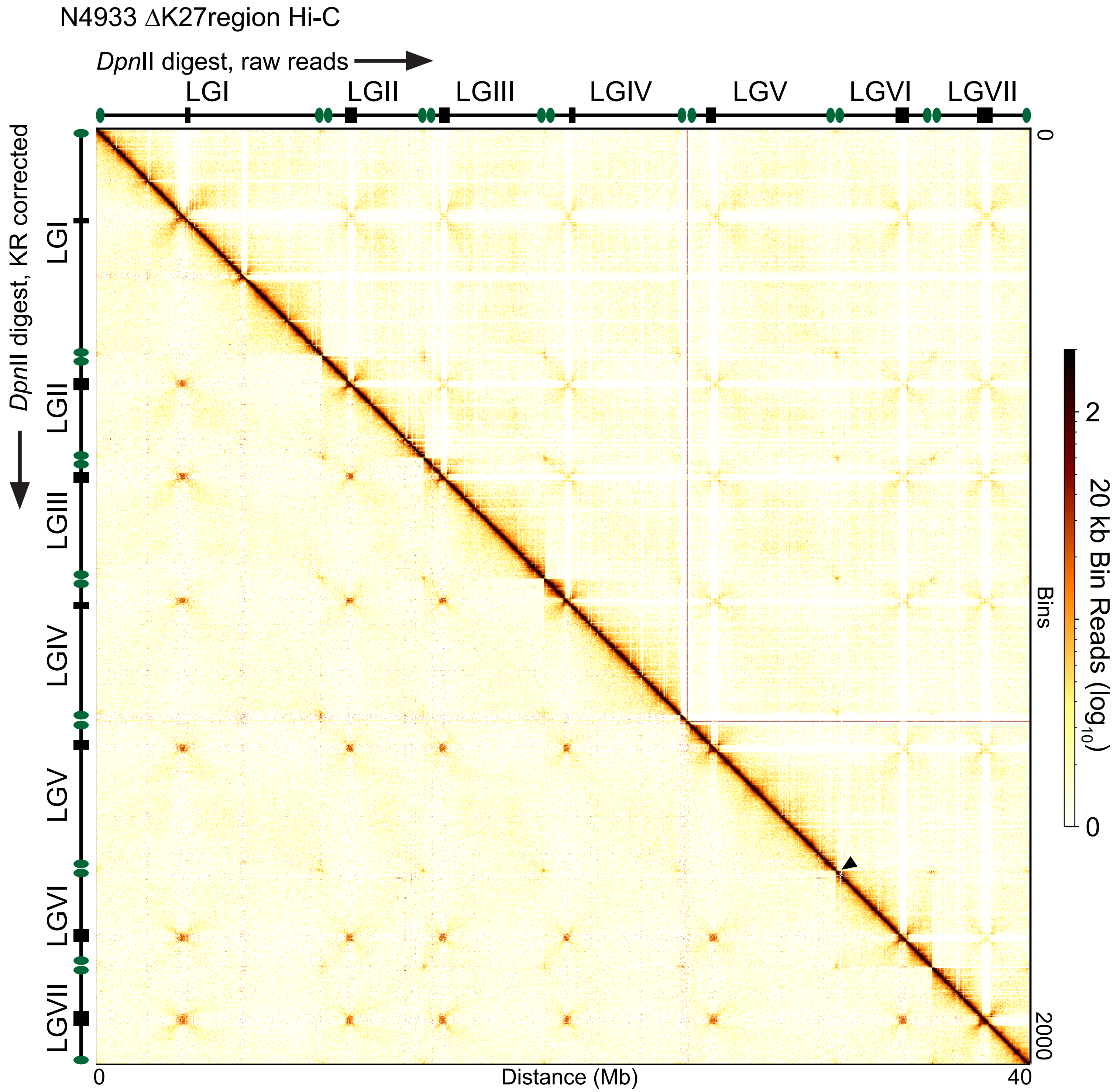


**Figure S2. Chromosome conformation of the whole genome of the N4933 ΔK27region strain using the restriction enzyme *Dpn*II, presented with either raw read counts or KR corrected counts.** Heatmap displaying *in situ* Hi-C contacts, either as (above diagonal) raw read counts or (below diagonal) KR-corrected counts for the N4933 ΔK27region strain across the entire *Neurospora crassa* genome at 20 kb bins. Black arrowhead indicates the site of the ΔK27region deletion.
