## Supplemental_Figure-3 for "A Constitutive Heterochromatic Region Shapes Genome Organization and Impacts Gene Expression in *Neurospora crassa*"

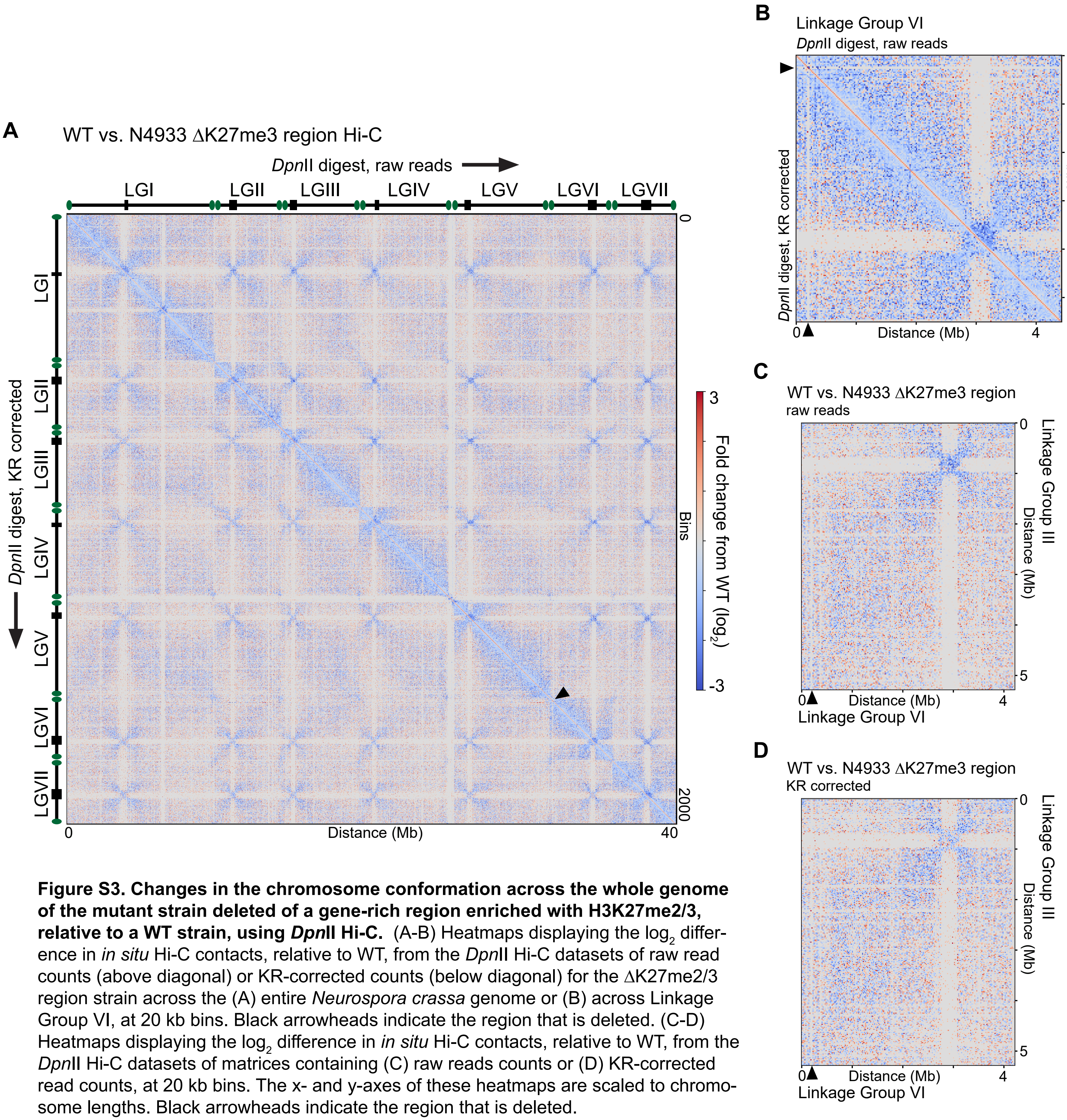


**Figure S3. Changes in the chromosome conformation across the whole genome of the mutant strain deleted of a gene-rich region enriched with H3K27me2/3, relative to a WT strain, using *Dpn*II Hi-C.** (A-B) Heatmaps displaying the $\log_2$ difference in *in situ* Hi-C contacts, relative to WT, from the *Dpn*II Hi-C datasets of raw read counts (above diagonal) or KR-corrected counts (below diagonal) for the ΔK27me2/3 region strain across the (A) entire *Neurospora crassa* genome or (B) across Linkage Group VI, at 20 kb bins. Black arrowheads indicate the region that is deleted. (C-D) Heatmaps displaying the $\log_2$ difference in *in situ* Hi-C contacts, relative to WT, from the *Dpn*II Hi-C datasets of matrices containing (C) raw reads counts or (D) KR-corrected read counts, at 20 kb bins. The x- and y-axes of these heatmaps are scaled to chromosome lengths. Black arrowheads indicate the region that is deleted.
