## Supplemental_Figure-4 for "A Constitutive Heterochromatic Region Shapes Genome Organization and Impacts Gene Expression in *Neurospora crassa*"

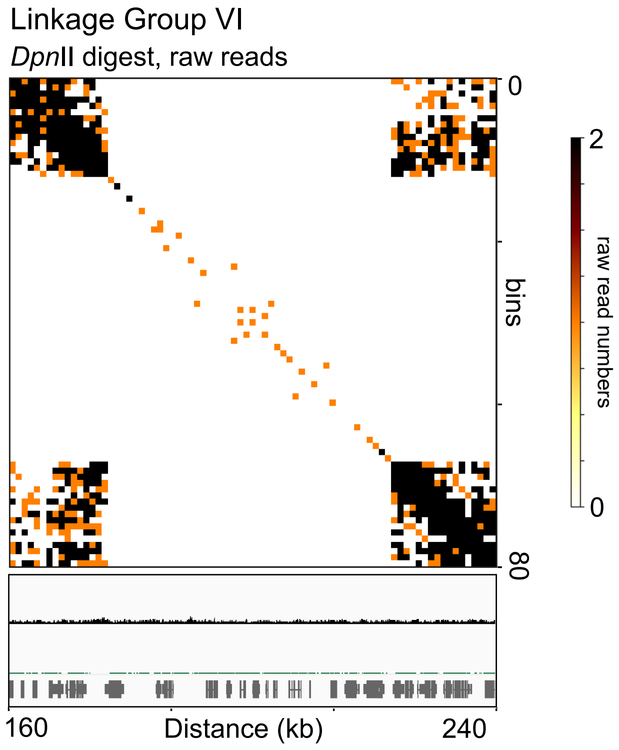


**Figure S4. Raw contact probability heatmaps of the ΔK27 region show small numbers of contacts prior to matrix correction.** Heatmaps displaying the numbers of raw contacts assigned to the ΔK27 region in a high-resolution (1kb) contact probability matrix. Scalebar (0 to 2 reads) to the right. IGV image of input DNA (black; SRR11806718) and H3K9me3 ChIP-seq data (green), with the former showing potential repetitive DNA sequences that can be amplified during library construction.
