## Supplemental_Figure-5 for "A Constitutive Heterochromatic Region Shapes Genome Organization and Impacts Gene Expression in *Neurospora crassa*"

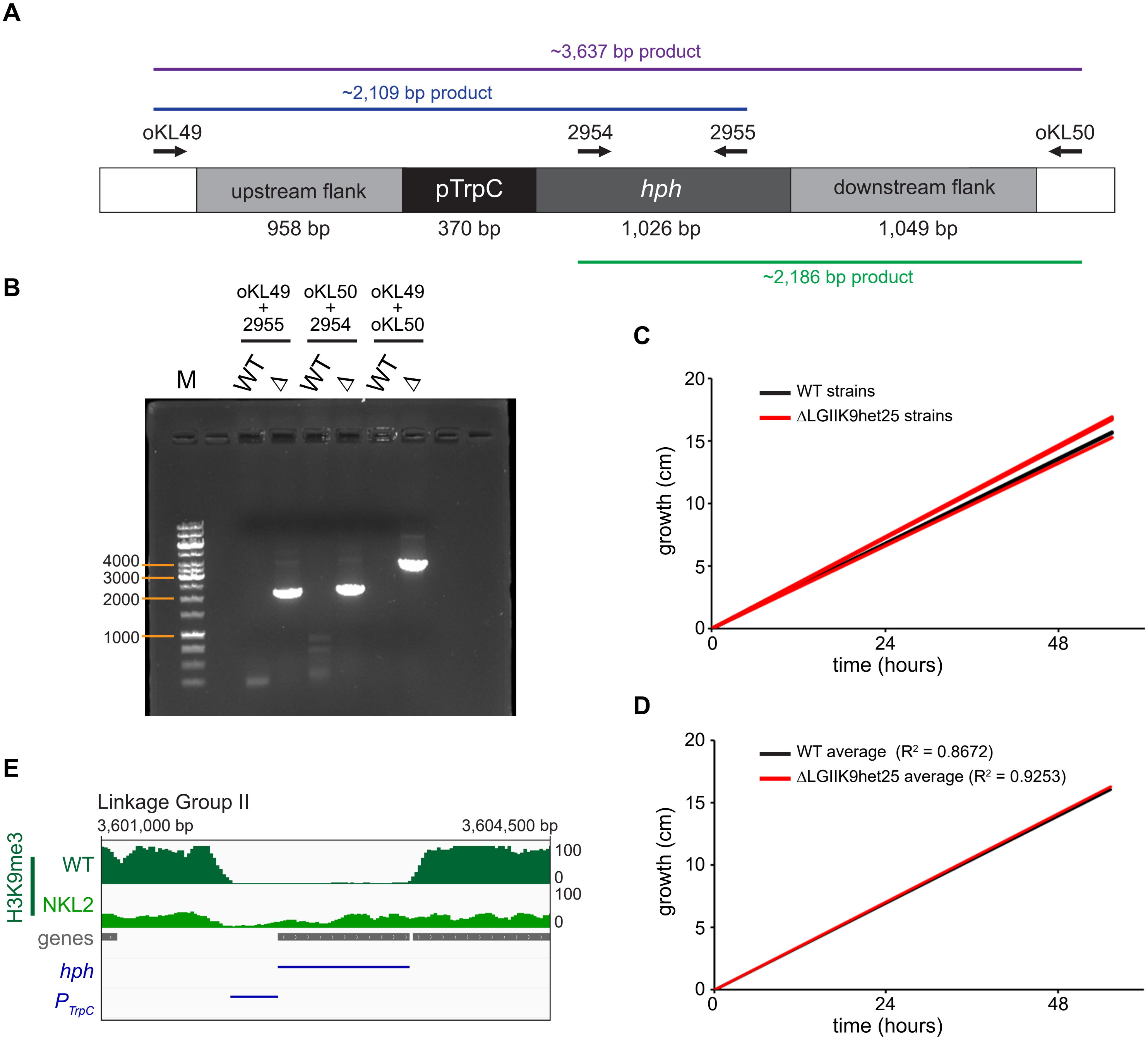


**Figure S5. Confirmation of the proper deletion of H3K9me3-marked heterochromatic region #25 on Linkage Group II and characterization of the resulting strain, NKL2 (ΔLGIIK9het25).** (A) Schematic of the insertion of the $P_{trpC}$*::hph* cassette into the H3K9me3-enriched AT-rich locus on Linkage Group II (LG II) from 3,602,033 bp to 3,712,857 bp, based on the *N. crassa* reference genome version 14. Arrows indicate relative positions of the oligonucleotides used for PCR in B, with names indicating these Primers. The gray boxes labeled "1 kb" indicate the DNA that flanks the heterochromatic region that was amplified for the initial stitching PCR reactions. Approximate sizes of the PCR products are shown in green, blue, and purple. Schematic is not to scale. This construct, removes 109,208 bp from the Neurospora genome (110,609 bp of AT-rich DNA deleted from the genome, but the $P_{trpC}$*::hph* cassette adds in 1,401 bp). (B) PCR reactions using wild type ("WT") genomic DNA or genomic DNA from a hygromycin resistant progeny ("Δ") isolated from a cross of the hygromycin resistant primary transformant (delta LGIIK9het25) with a wild type strain. The hygromycin resistant primary transformant was isolated following transformation of the split marker constructs. This cross progeny xKL9AR-17 was used to establish the NKL2 strain. M = GeneRuler 1 kb DNA ladder (ThermoFisher Scientific). (C-D) Race tube growth analysis of strains with a WT or NKL2 (ΔLGIIK9het25) genotype, either in (C) triplicate or D) the average of the triplicate growth, plotting centimeters grown over hours of time. Trendlines for growth are shown with $R^2$ correlation values indicated for the triplicate experiments in (D). (E) Integrative Genome Viewer (IGV) tracks of H3K9me3 data from WT and NKL2 strains mapped to the NKL2 genome at the site of the $P_{trpC}$*::hph* cassette insertion. The y-axis scale is set from 0 to 100, which is enhancing the background signal from these ChIP-seq experiments. Motifs that indicate the $P_{trpC}$ and *hph* DNA sequences shown below.
