## Supplemental_Figure-6 for "A Constitutive Heterochromatic Region Shapes Genome Organization and Impacts Gene Expression in *Neurospora crassa*"

**A**

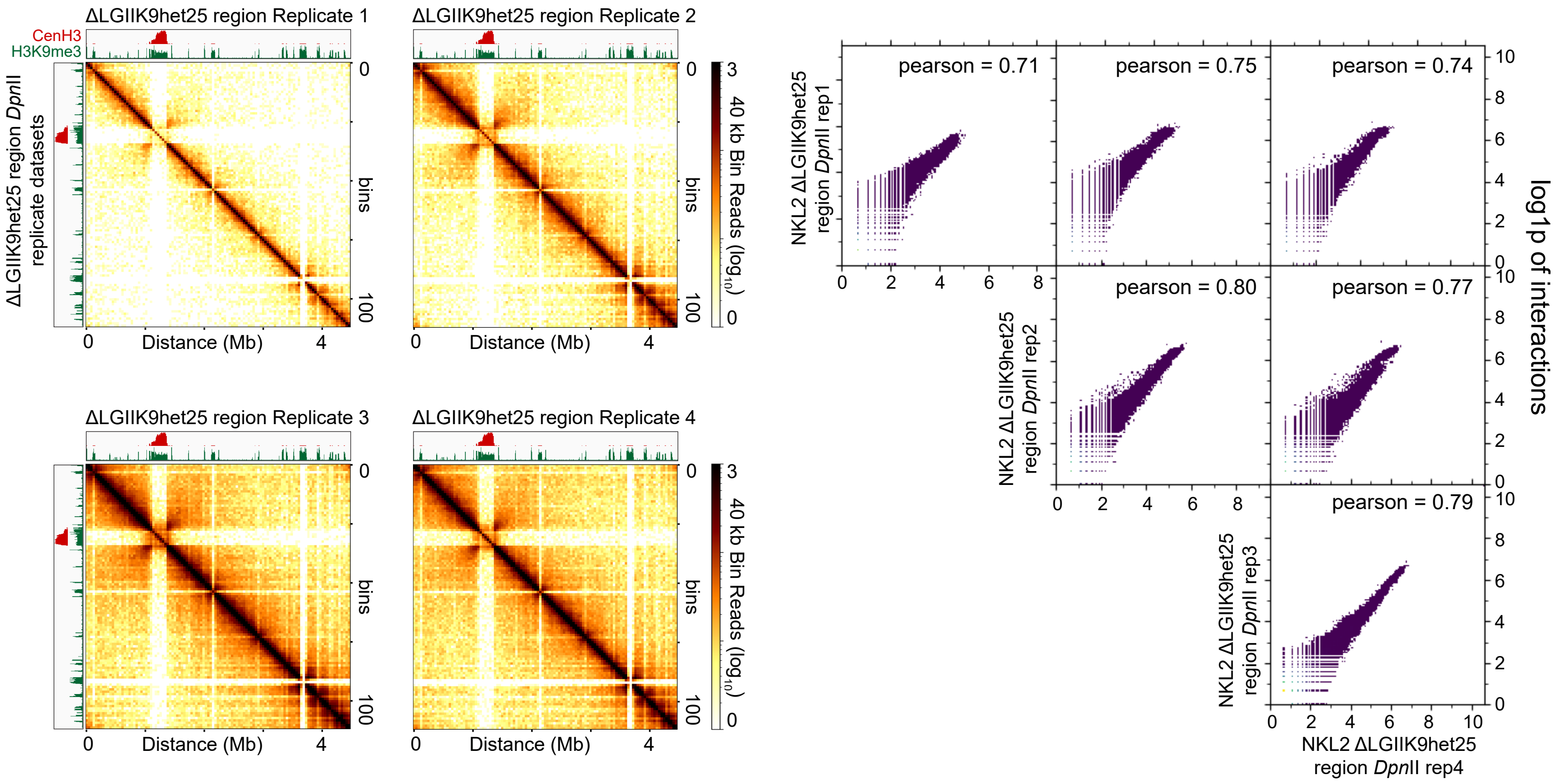


**B**

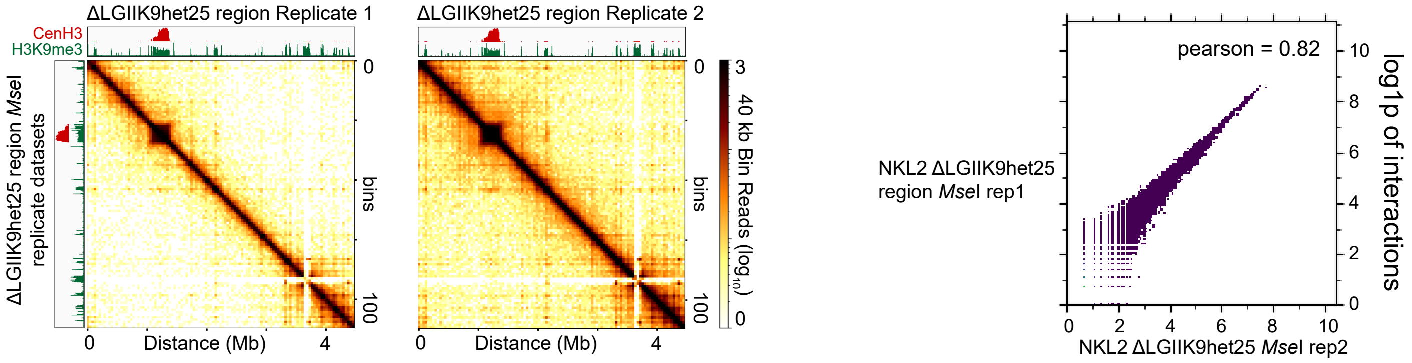


**Figure S6. Replicate analysis of *in situ* Hi-C genome organization datasets, using the *Mse*I restriction enzyme, of the ΔLGIIK9het25 strain of *Neurospora crassa*.** (A-B) Heatmaps and scatterplots of NKL2 ΔLGIIK9het25 replicates digested with (A) *Dpn*II or (B) *Mse*I. In each panel, the left images show the raw count Hi-C heatmaps of genomic interactions across LG II of each *in situ* Hi-C replicate dataset at 40 kb bin resolution, while the right image shows a scatter plot comparing interactions between each NKL2 ΔLGIIK9het25 replicate matrix at 20kb bin resolution. For scatterplots, log1p values of interactions presented and Pearson correlation values of each binary comparison shown at the upper right; Scatterplot image produced by the hicCorrelate program in hicExplorer.
