## Supplemental_Figure-10 for "A Constitutive Heterochromatic Region Shapes Genome Organization and Impacts Gene Expression in *Neurospora crassa*"

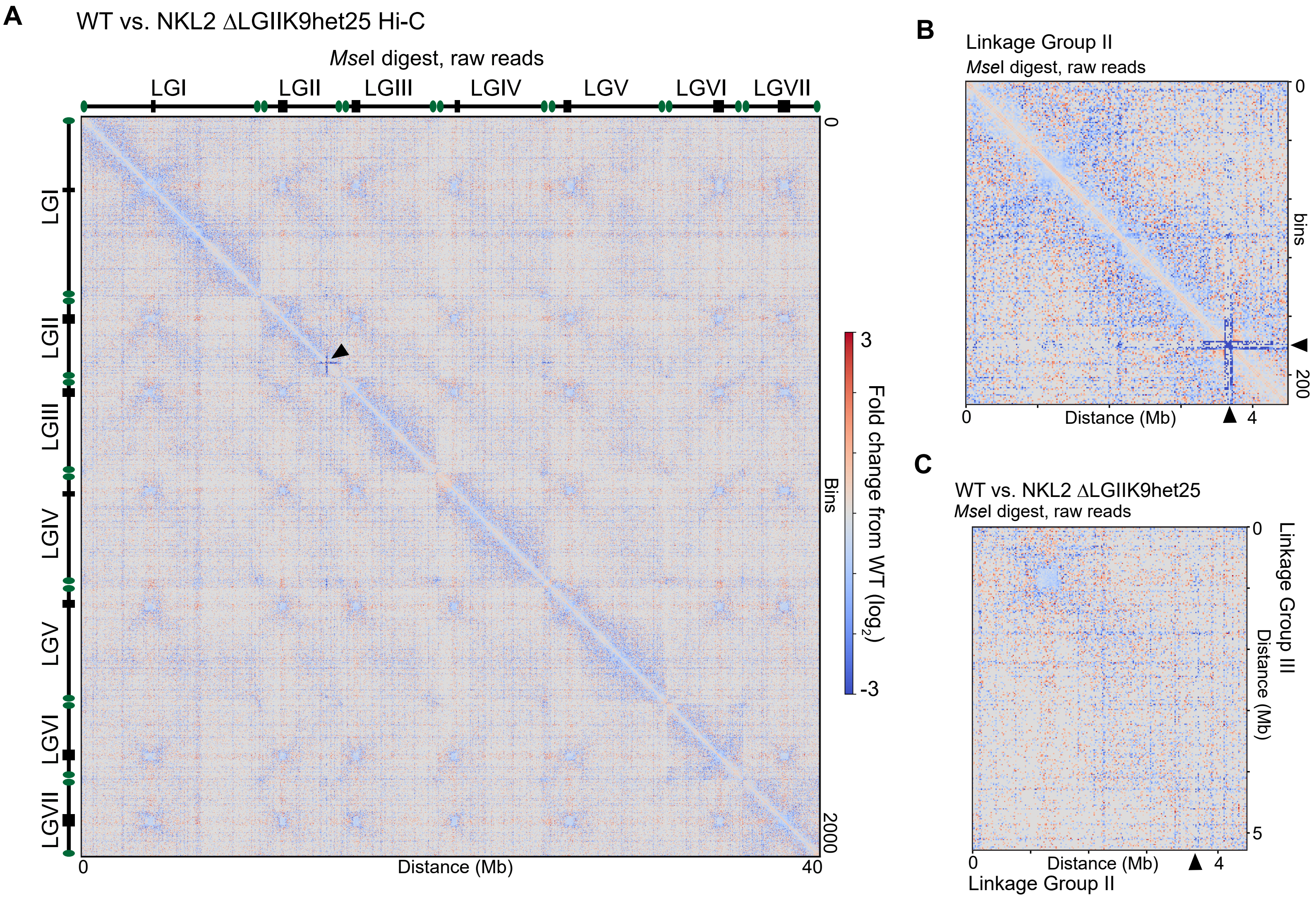


**Figure S10. Changes in the chromosome conformation across the whole genome of the mutant strain deleted of a gene-poor, AT-rich region enriched with H3K9me3, relative to a WT strain, using *Mse*I Hi-C.** (A-B) Heatmaps displaying the $log_2$ difference in *in situ* Hi-C contacts, relative to WT, from the *Mse*I Hi-C datasets of raw read counts for the ΔLGIIK9het25 region strain (NKL2) across the (A) entire *Neurospora crassa* genome or (B) across Linkage Group II, at 20 kb bins. Black arrowheads indicate the region that is deleted. (C) Heatmaps displaying the $log_2$ difference in *in situ* Hi-C contacts, relative to WT, from the *Mse*I Hi-C datasets of matrices containing raw reads counts at 20 kb bins. The x- and y-axes of this heatmap is scaled to chromosome lengths. Black arrowheads indicate the region that is deleted.
