## Supplemental_Figure-12 for "A Constitutive Heterochromatic Region Shapes Genome Organization and Impacts Gene Expression in *Neurospora crassa*"

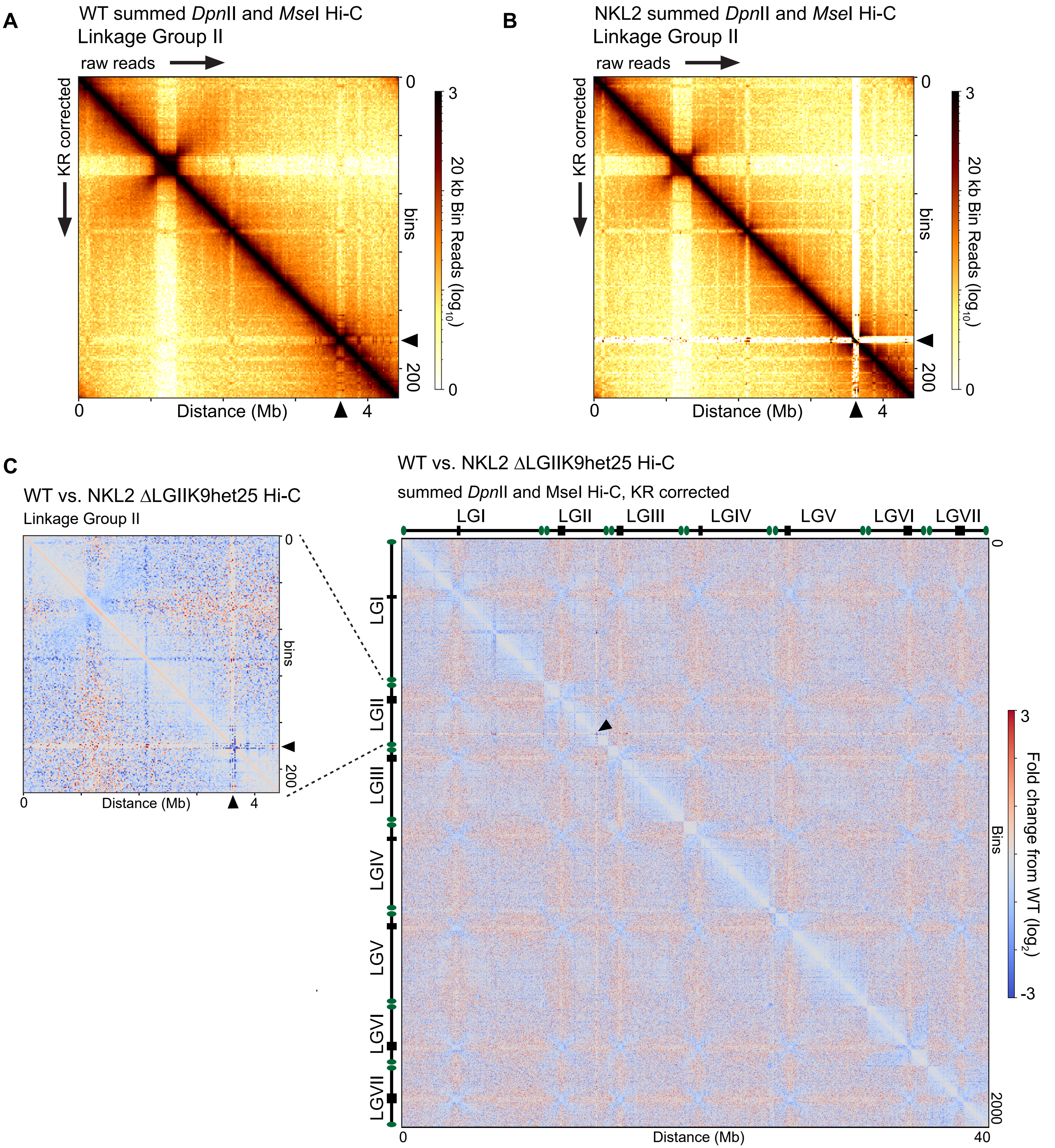


**Figure S12. Hi-C matrices in which *Dpn*II and *Mse*I *in situ* Hi-C datasets from WT and ΔLGIIK9het25 strains are independently summed show changes in contact probability across the Neurospora genome.** (A-B) Contact probability heatmaps displaying the $\log_{10}$ read count for *Dpn*II and *Mse*I summed Hi-C datasets of raw read counts (above diagonal) or KR-corrected counts (below diagonal) for the (A) WT strain or (B) NKL2 ΔLGIIK9het25 strain across Linkage Group II, at 20 kb bins. Black arrowheads indicate the region that is deleted. (C) Heatmaps displaying the $\log_2$ difference in *in situ* Hi-C contacts in a NKL2 ΔLGIIK9het25 strain, relative to WT, of *Dpn*II and *Mse*I summed Hi-C datasets of matrices containing KR-corrected read counts, at 20 kb bins across the entire Neurospora genome or LG II (zoomed in region). Black arrowheads indicate the region that is deleted.
