## Supplemental_Figure-13 for "A Constitutive Heterochromatic Region Shapes Genome Organization and Impacts Gene Expression in *Neurospora crassa*"

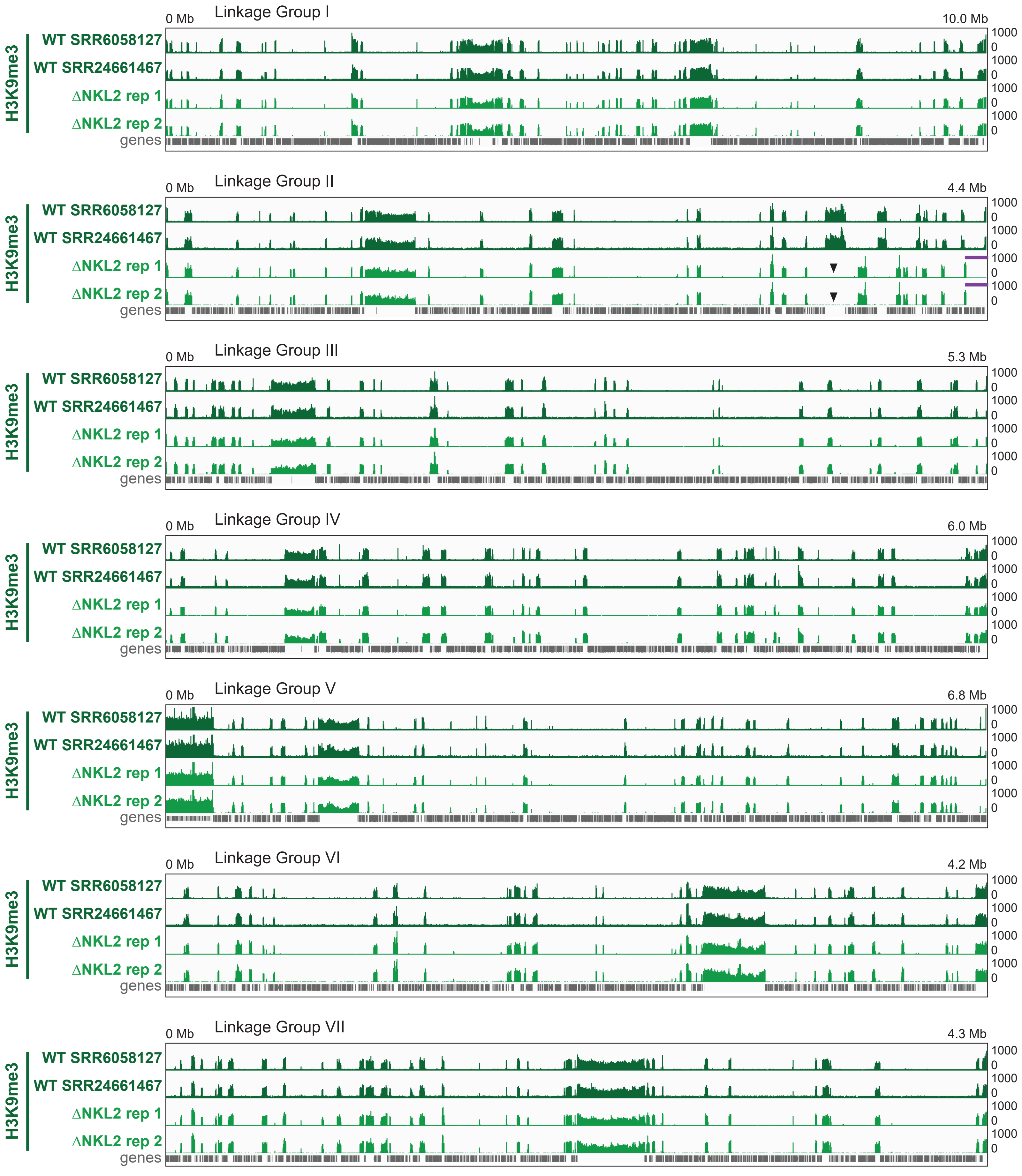


**Figure S13. Replicates of H3K9me3 ChIP-seq in NKL2 (ΔLGIIK9het25) strains are reproducible.** Integrative Genomics Viewer (IGV) images of tracks of two previously published WT H3K9me3 ChIP-seq replicate datasets (SRR6058127, top; SRR24661467, bottom; numbers from the NIH Sequence Read Archive), and two replicate H3K9me3 ChIP-seq datasets of the NKL2 strain (ΔLGIIK9het25) generated here. Genes shown in gray. ChIP-seq enrichment value noted at the right, while chromosome distances presented at the top. The site of the heterochromatic region LGIIK9het25 deleted from Linkage Group II in NKL2 strains shown by arrowheads, while the purple line indicates the region of Linkage Group II that has no ChIP-seq signal due to the loss of the DNA ΔLGIIK9het25 deletion shifting other ChIP-seq reads. All ChIP-seq reads were mapped to the WT “nc14” reference genome.
