## Supplemental_Figure-14 for "A Constitutive Heterochromatic Region Shapes Genome Organization and Impacts Gene Expression in *Neurospora crassa*"

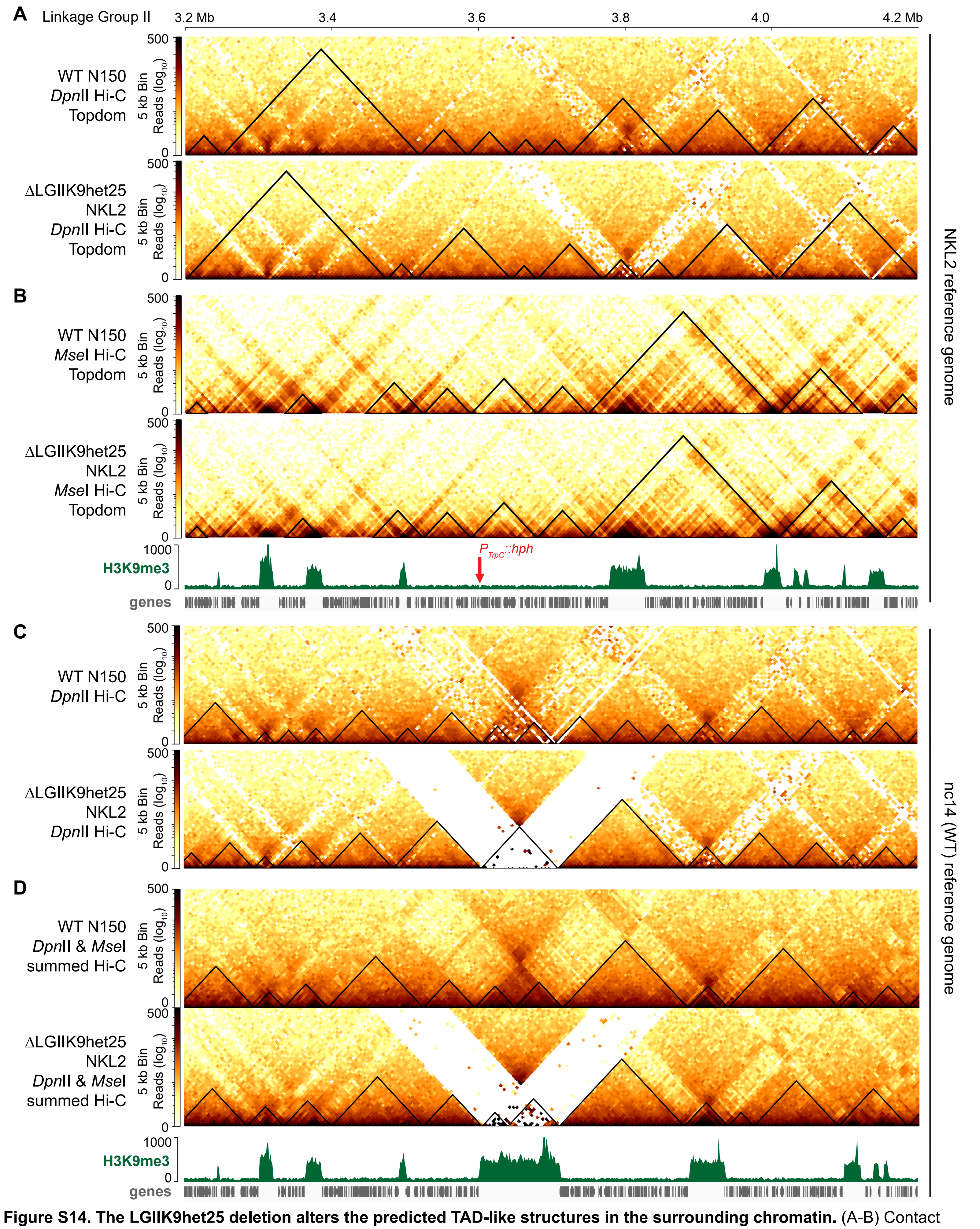


**Figure S14. The LGIIK9het25 deletion alters the predicted TAD-like structures in the surrounding chromatin.** (A-B) Contact probability heatmaps of (A) *Dpn*II (KR corrected) or (B) *Mse*I (raw) *in situ* Hi-C datasets of WT and ΔLGIIK9het25 strains at 5 kb bin resolution mapped to the NKL2 ΔLGIIK9het25 reference genome. Wild type H3K9me3 ChIP-seq (green) and gene (gray) tracks shown below. The contact probability scalebar is shown to the left of each heatmap. Black triangles show predicted TAD-like structures, as calculated by the program Topdom. The red arrow shows the position of the $P_{trpC}$*::hph* gene in the NKL2 reference genome. (C-D) Contact probability heatmaps of (A) *Dpn*II (KR corrected) or (B) summed *Dpn*II and *Mse*I (KR corrected) *in situ* Hi-C datasets of WT and ΔLGIIK9het25 strains at 5 kb bin resolution mapped to the nc14 (WT) reference genome. TAD-like structures are predicted by the program hicFindTADs. Wild type H3K9me3 ChIP-seq (green) and gene (gray) tracks shown below.
