## Supplemental_Figure-15 for "A Constitutive Heterochromatic Region Shapes Genome Organization and Impacts Gene Expression in *Neurospora crassa*"

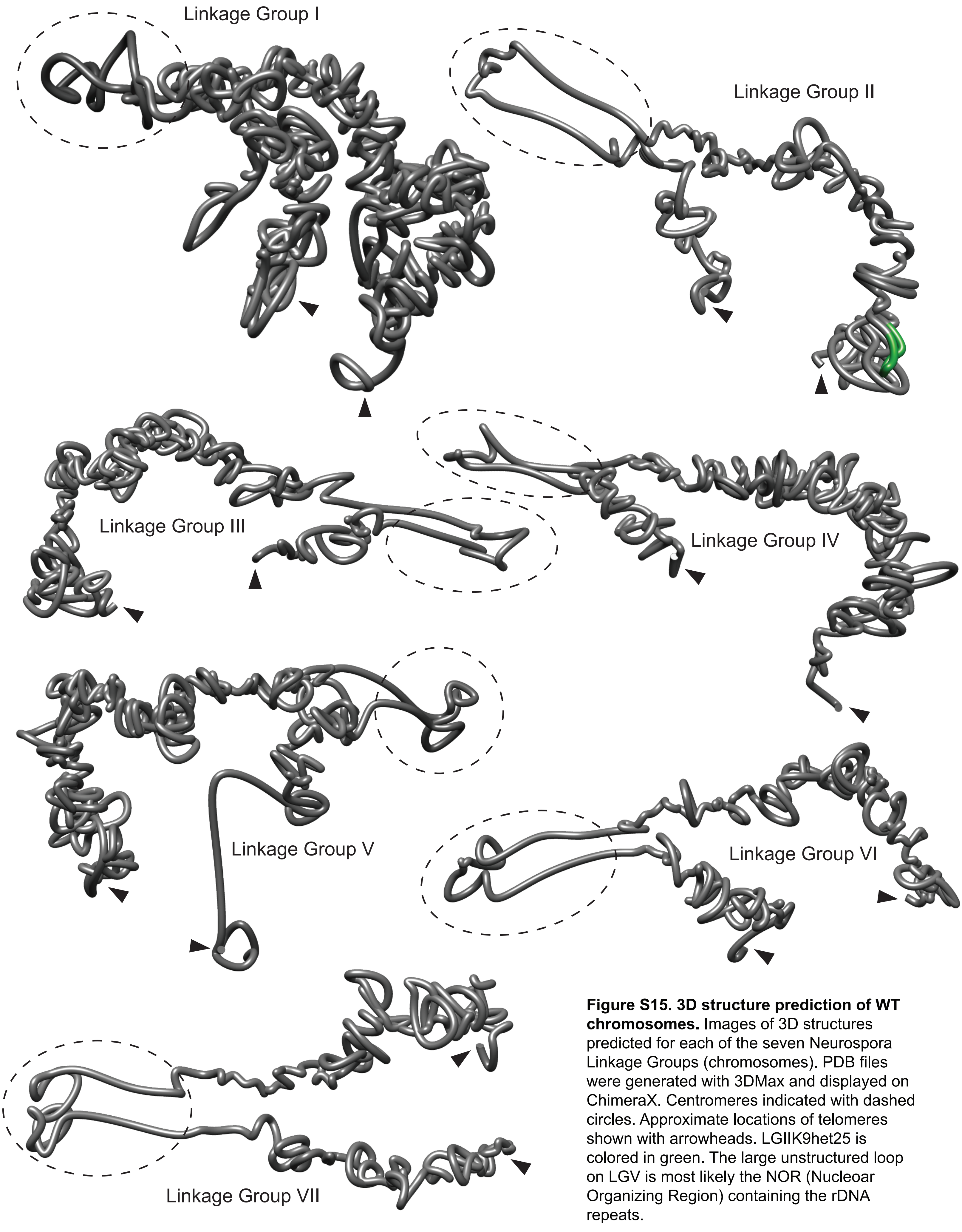


**Figure S15. 3D structure prediction of WT chromosomes.** Images of 3D structures predicted for each of the seven Neurospora Linkage Groups (chromosomes). PDB files were generated with 3DMax and displayed on ChimeraX. Centromeres indicated with dashed circles. Approximate locations of telomeres shown with arrowheads. LGIIK9het25 is colored in green. The large unstructured loop on LGV is most likely the NOR (Nucleoar Organizing Region) containing the rDNA repeats.
