## Supplemental_Figure-17 for "A Constitutive Heterochromatic Region Shapes Genome Organization and Impacts Gene Expression in *Neurospora crassa*"

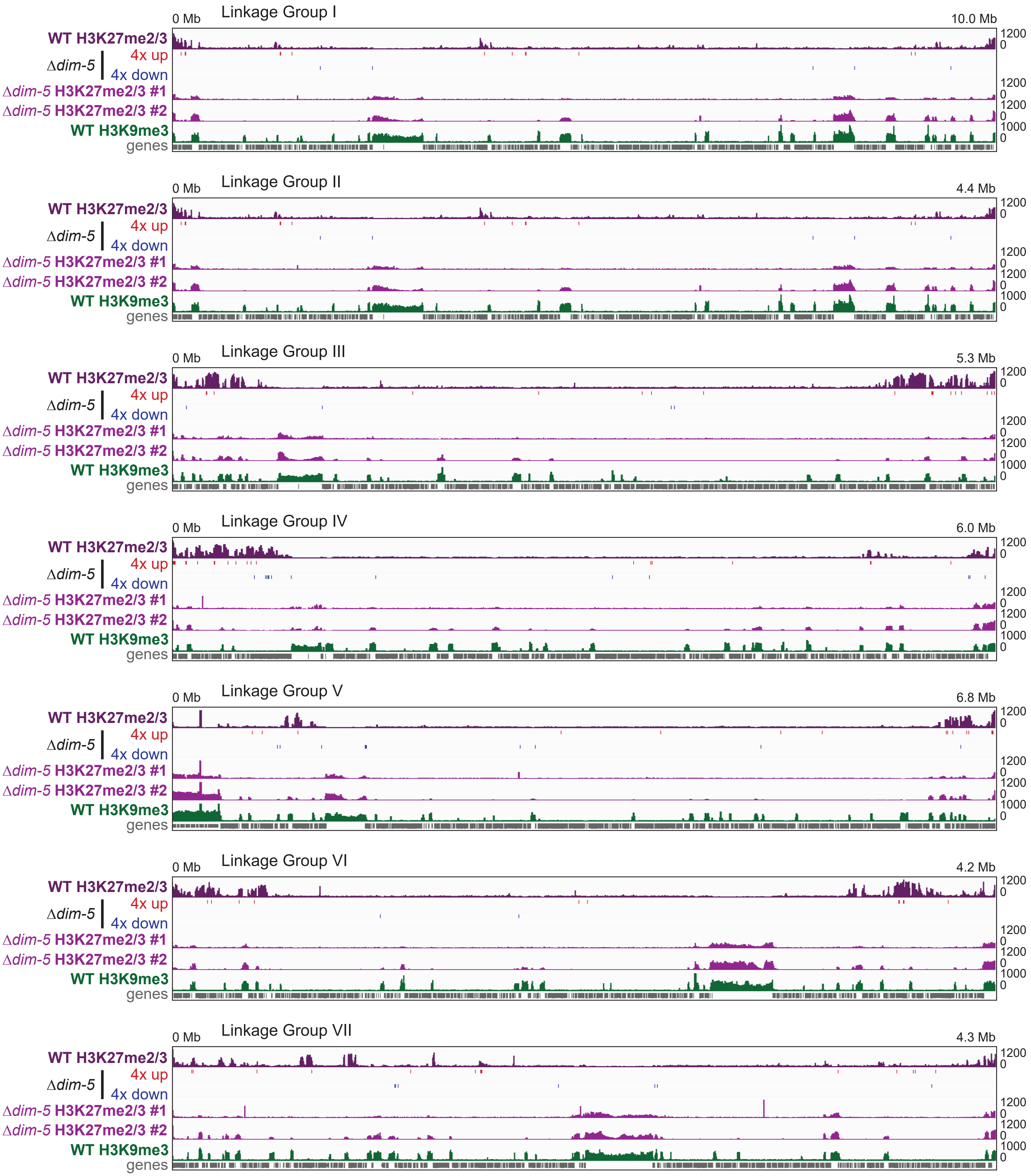


**Figure S17. Altered gene expression in a Δ*dim-5* strain, relative to a WT strain, might be due to relocalization of facultative heterochromatin.** Integrative Genomics Viewer (IGV) images of tracks of the WT H3K27me2/3 ChIP-seq, genes with altered expression in the Δ*dim-5* (N3944) strain, relative to a WT (N150) strain, previously published Δ*dim-5* H3K27me2/3 ChIP-seq datasets (#1 = SRR2026374; #2 = SRR2036169; SRR numbers from the NIH Sequence Read Archive), and WT H3K9me3 ChIP-seq datasets. All differentially expressed genes have a $\log_2 > 4.0$ and adjusted p-value < 0.001. Genes shown in gray. ChIP-seq enrichment value noted at the right, while chromosome distances presented at the top.
