## Supplemental_Figure-18 for "A Constitutive Heterochromatic Region Shapes Genome Organization and Impacts Gene Expression in *Neurospora crassa*"

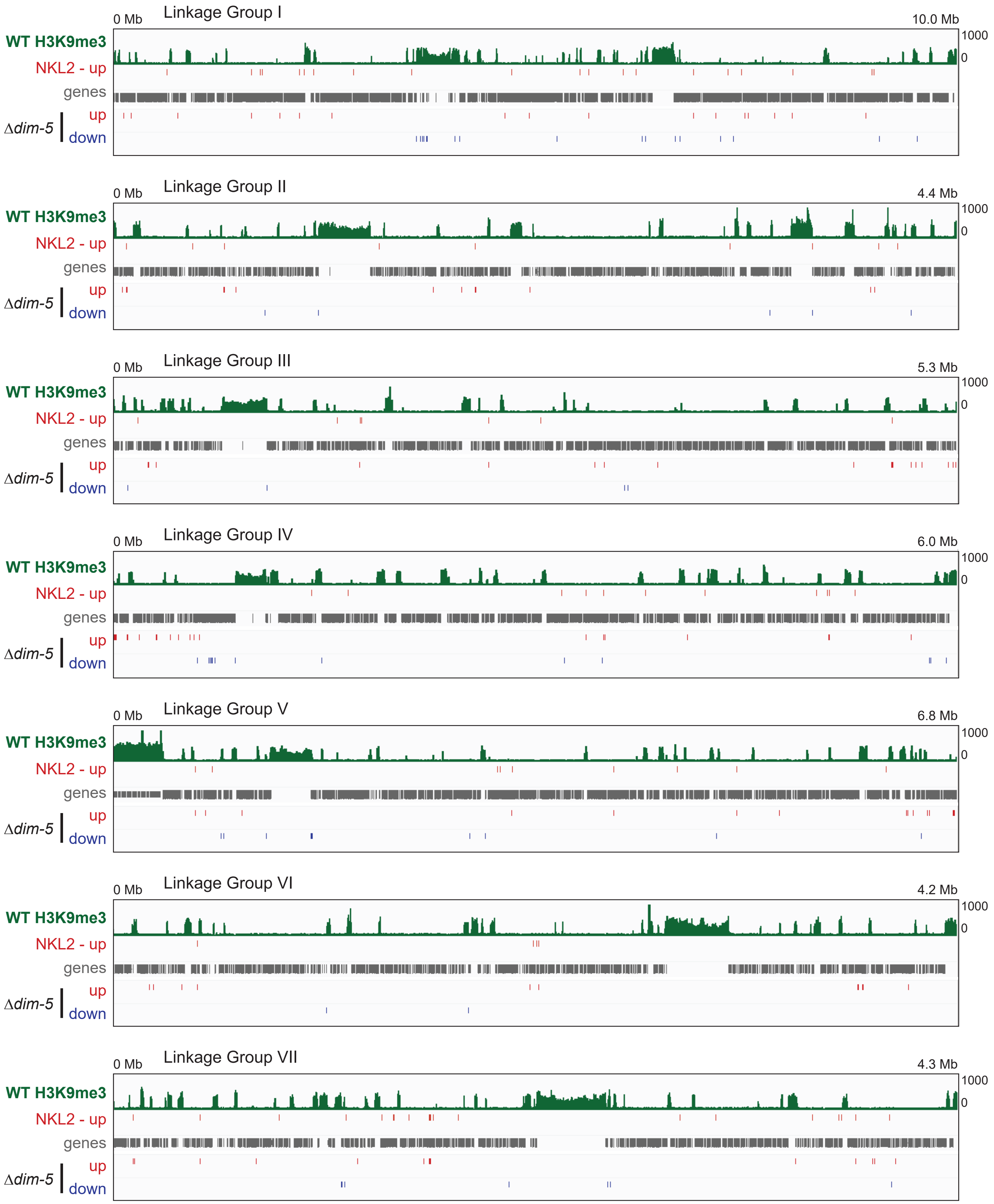


**Figure S18. Differential Expression analysis of genes with altered expression in NKL2 (ΔLGIIK9het25) and Δ*dim-5* strains, relative to a WT strain, show altered expression across the Neurospora genome.** Integrative Genomics Viewer (IGV) images of the WT H3K9me3 (top) and genes with altered expression in either NKL2 or Δ*dim-5* (N3944) strains, relative to a WT (N150) strain. All differentially expressed genes have a $\log_2 > 3.0$ ("up" genes) or $< -3.0$ ("down" genes) and adjusted p-value < 0.001. Genes shown in gray. Enrichment value of H3K9me3 ChIP-seq noted at the right, while chromosome distances presented at the top.
