## Supplemental_Figure-19 for "A Constitutive Heterochromatic Region Shapes Genome Organization and Impacts Gene Expression in *Neurospora crassa*"

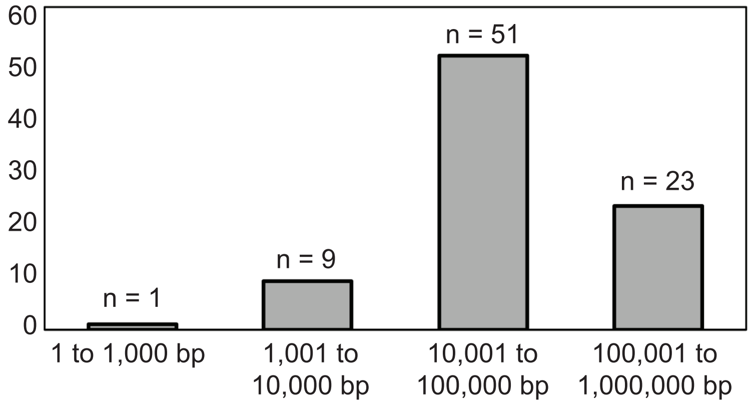


**Figure S19. Differentially Expressed Genes are distant from H3K9me3-marked heterochromatic regions.** The distance from the Transcription Start Site of each Differentially Expressed Gene (DEG) in an NKL2 ΔLGIIK9het25 strain, relative to a WT strain, to the nearest H3K9me3 boundary of a constitutive heterochromatic region. The genes are grouped into four distance categories, with the number of genes in each group listed at the top of the bar. The average gene distances in each group are: from 1 to 1,000 bp = 95 bp; from 1,001 to 10,000 bp = 6397 bp; from 10,001 to 100,000 bp = 38,255 bp; from 100,001 to 1,000,000 bp = 193,001 bp.
