## Supplemental_Figure-21 for "A Constitutive Heterochromatic Region Shapes Genome Organization and Impacts Gene Expression in *Neurospora crassa*"

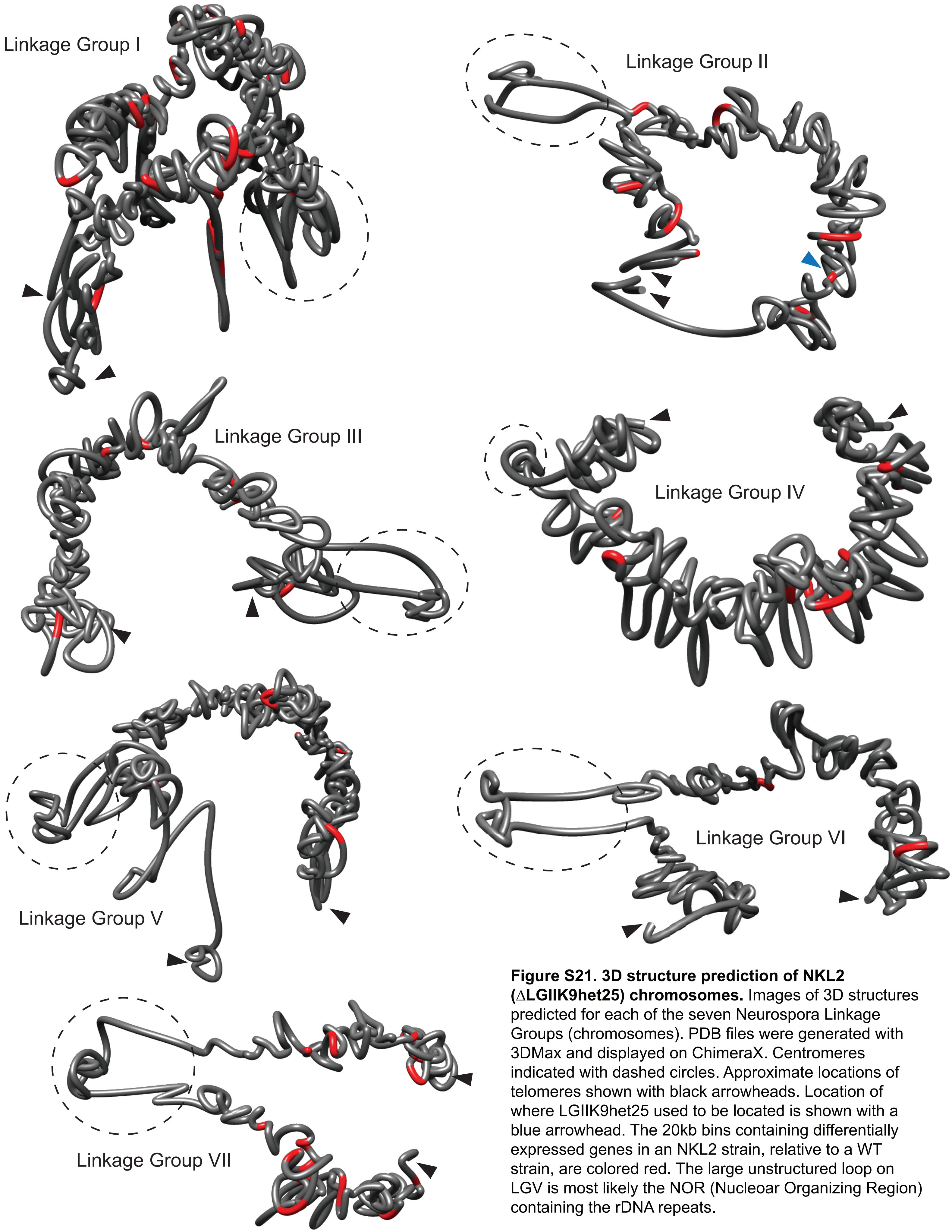


**Figure S21. 3D structure prediction of NKL2 (∆LGIIK9het25) chromosomes.** Images of 3D structures predicted for each of the seven Neurospora Linkage Groups (chromosomes). PDB files were generated with 3DMax and displayed on ChimeraX. Centromeres indicated with dashed circles. Approximate locations of telomeres shown with black arrowheads. Location of where LGIIK9het25 used to be located is shown with a blue arrowhead. The 20kb bins containing differentially expressed genes in an NKL2 strain, relative to a WT strain, are colored red. The large unstructured loop on LGV is most likely the NOR (Nucleoar Organizing Region) containing the rDNA repeats.
