## Supplemental_Figure-22 for "A Constitutive Heterochromatic Region Shapes Genome Organization and Impacts Gene Expression in *Neurospora crassa*"

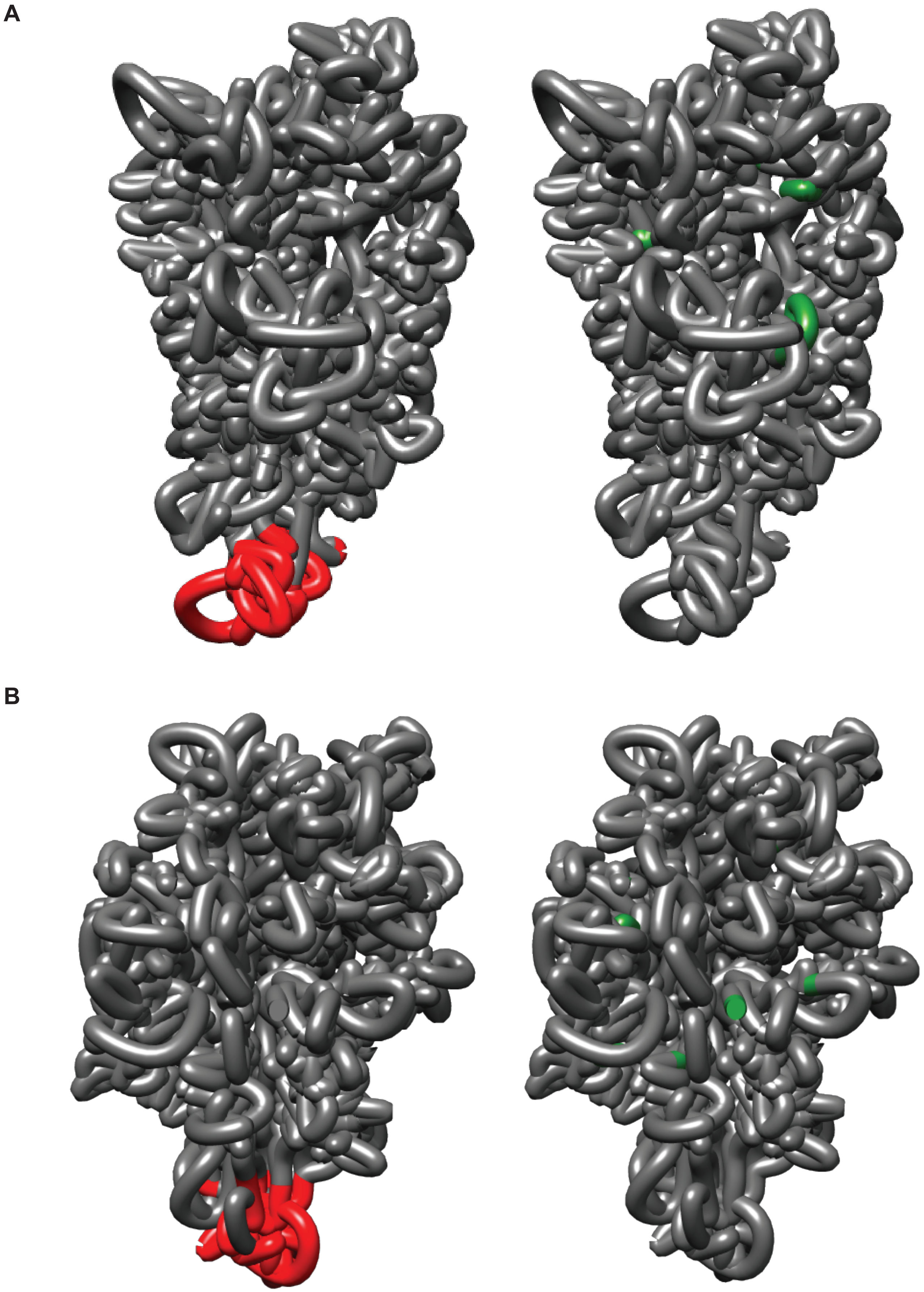


**Figure S22. 3D structure prediction of the WT and NKL2 genomes show biologically relevant interactions consistent with a Rabl chromosome conformation.** (A-B) Images of 3D structures predicted for the (A) WT and (B) NKL2 genomes at 20 kb bin resolution indicating (left) centromeres or (right) telomeres. PDB files were generated with LorDG and displayed on ChimeraX. Centromeric bins, as defined by enrichment of CenH3::mCherry (Galazka et al., 2016), are colored red, while chromosome ends that indicate telomeres are colored green.
