## Supplemental_Table-1 for "A Constitutive Heterochromatic Region Shapes Genome Organization and Impacts Gene Expression in *Neurospora crassa*"

Supplementary Table S1 – oligonucleotides

Reckard et al.,

A Constitutive Heterochromatic Region Shapes Genome Organization and Impacts Gene Expression in *Neurospora crassa*

| ***Oligo name*** | ***Target*** | ***direction*** | ***sequence*** |
| --- | --- | --- | --- |
| 2954 | *hph* | reverse | 5’ tcgcctcgctccagtcaatgacc |
| 2955 | *hph* | forward | 5’ aaaaagcctgaactcaccgcgacg |
| oKL14 | LGIIK9het25 upstream sequence | forward | 5’ TCCTGAGCAGTGAACAAGCC |
| oKL15 | LGIIK9het25 upstream sequence | reverse | 5’ ttctgtcgacGCTGTGGACCTTATCGAGCC |
| oKL18 | LGIIK9het25 downstream sequence | forward | 5' aatagagtagCCGCTTACCAGGTAACATCCG |
| oKL19 | LGIIK9het25 downstream sequence | reverse | 5’ AGGCTCGAGCAATTTGTCGC |
| oKL49 | LGIIK9het25 upstream sequence – beyond boundary | forward | 5’ AAGAGACGAGGAGAACTGGCC |
| oKL50 | LGIIK9het25 downstream sequence – beyond boundary | reverse | 5’ TTGGACAGCATCTTCGACGG |
| oKL55 | *mus-52* | forward | 5’GGTTAGTGGGAAGTGTCCGC |
| oKL56 | *mus-52* | reverse | 5’GGTTTGGATCATTCTGAGGGC |
